# Single cell transcriptomic and spatial landscapes of the developing human pancreas

**DOI:** 10.1101/2022.02.04.478971

**Authors:** Oladapo E. Olaniru, Ulrich Kadolsky, Shichina Kannambath, Heli Vaikkinen, Kathy Fung, Pawan Dhami, Shanta J. Persaud

**Affiliations:** Department of Diabetes, School of Cardiovascular and Metabolic Medicine & Sciences, King’s College London, Guy’s Campus, London SE1 1UL, UK; Genomics Research Platform and Single Cell Laboratory, Biomedical Research Centre, Guy’s and St. Thomas’ NHS Trust, London, UK; Genomics WA, University of Western Australia, Harry Perkins Institute of Medical Research and Telethon Kids Institute QEII Campus, Nedlands, Perth, WA 6009, Australia

**Keywords:** human fetal pancreas, scRNA-seq, Visium, spatial transcriptomics, beta cell development

## Abstract

The progress made in directed differentiation of stem cells has shown that understanding human pancreas development can provide cues for generating unlimited amounts of insulin-producing beta cells for transplantation therapy in diabetes. However, current differentiation protocols have not been successful in reproducibly generating functional human beta cells in vitro, partly due to incomplete understanding of human pancreas development. Here, we present detailed transcriptomic analysis of the various cell types of the developing human pancreas, including their spatial gene patterns. We integrated single cell RNA sequencing with spatial transcriptomics at multiple developmental timepoints and revealed distinct temporal-spatial gene cascades in the developing human pancreas. Cell trajectory inference identified endocrine progenitor populations and novel branch-specific genes as the progenitors differentiate towards alpha or beta cells, indicating that transcriptional maturation occurred over this developmental timeframe. Spatial differentiation trajectories indicated that immature Schwann cells are spatially co-located with endocrine progenitors and contribute to beta cell maturation via the L1CAM-EPHB2 pathway. Our integrated approach enabled us to identify heterogeneity and multiple lineage dynamics within the mesenchyme, showing that it contributed to the exocrine acinar cell state. Finally, we have generated an interactive web resource for interrogating human pancreas development for the research community.

## INTRODUCTION

The pancreas is a multicellular organ composed of exocrine and endocrine compartments. The exocrine pancreas contains acinar and ductal cells that secrete digestive juices, while the endocrine pancreas contains alpha, beta, delta, epsilon and pancreatic polypeptide cells that co-operatively regulate glucose homeostasis. Despite the critical roles of the pancreas in nutrient digestion and glucose homeostasis, the dysfunction of which results in pancreatitis, pancreatic cancers and diabetes affecting more than half a billion people worldwide (Bray et al., 2018; Ouyang et al., 2020; Saeedi et al., 2019), the mechanisms underlying how the individual cell types develop in humans remain unclear.

Understanding pancreas developmental trajectories can provide essential knowledge for generating unlimited amounts of insulin-producing beta cells, for example, from stem cells for cell replacement therapies of type 1 diabetes, and much work has been expended in recent years to define effective differentiation protocols (D’Amour et al., 2006; Pagliuca et al., 2014; Russ et al., 2015; Veres et al., 2019). However, current differentiation strategies, which are mainly based on recapitulating gene cascades identified in mouse pancreas development, do not reproducibly produce fully functional human beta cells in vitro (Nair et al., 2019; Rezania et al., 2014; Russ et al., 2015; Zhu et al., 2016). This is not surprising as human islet development occurs over a longer time-span than in mice and inter-species differences have been reported, such as in the islet cytoarchitecture, and the presence of a single wave of endocrine differentiation and delayed expression of key differentiation genes in humans (Nair and Hebrok, 2015; Villasenor et al., 2008). A clearer understanding of human pancreatic endocrinogenesis is therefore required, and some progress has recently been made in this through single cell RNA sequencing (scRNA-seq). Thus, different progenitor populations have been identified in the very early stages of pancreas development (7 and 10 post conception week; PCW) (Gonçalves et al., 2021) and significant differences in lineage differentiation between developing mouse and human pancreas have recently been identified (Yu et al., 2021).

However, while scRNA-seq provides a snapshot of gene expression profiles at an unprecedented scale, gene information in relation to spatial cell context is lost since tissue dissociation is required. Spatial transcriptomics gives positional gene patterns and provides spatial attributes of cells within the tissue context. In this study we have therefore utilised both high throughput scRNA-seq and spatial transcriptomics of human fetal pancreases at multiple developmental stages followed by data integration to define the cellular heterogeneity and spatial developmental landscape of human pancreas develoment. This approach has allowed us to chararacterise and spatially resolve multiple human pancreatic cell populations at different developmental stages, including their cell-cell interactions, and we have identified novel gene candidates that regulate progenitor cell differentiation. By estimating pairwise similarity in transcriptional profiles among cells, we have uncovered spatial differentiation trajectories in situ, which enabled us to identify, for the first time, the importance of Schwann cells and mesenchymal cells in the differentiation of human endocrine progenitors and acinar cells, respectively.

## RESULTS

### scRNA-seq analysis of whole human pancreases at 12-20 PCW

We combined scRNA-seq with 10x Visium spatial transcriptomics to systematically characterise the developing landscape of the human pancreas, as indicated in Fig 1A. For scRNA-seq, whole pancreases from 12 individual embryos spanning 7 developmental time points (12, 13, 14, 15, 18, 19 and 20 PCW) were dissociated, and live-sorted single cells were multiplexed with antibody-oligonucleotide conjugates that bind to the ubiquitous surface markers β2M and CD298 (Stoeckius et al., 2018), which are expressed by the developing human pancreas (Suppl. Fig 1A-1F). We sequenced the cells using the 10x Chromium protocol and retained 24,080 high quality cells following stringent filtering for downstream analysis (12pcw: 4099, 13pcw: 7168, 14pcw: 3530, 15pcw: 1664, 18pcw: 2171, 19pcw: 3827, 20pcw: 1621; Suppl. Fig 1G-1I). Unsupervised clustering of our scRNA-seq data revealed distinct populations of acinar, ductal, endocrine, endothelial, erythroblast, immune, mesenchymal and Schwann cells (Fig 1B & 1C), which we identified by differential expression of established markers (Fig 1D & 1E). For example, acinar cells were identified by expression of CPA1, endocrine cells by CHGA, mesenchyme by COL3A1, immune cells by RAC2, ductal cells by CFTR, endothelial cells by ADGRL4, erythroblasts by HBB and Schwann clusters by CRYAB expression (Fig 1 D & 1E).

**Fig 1:**
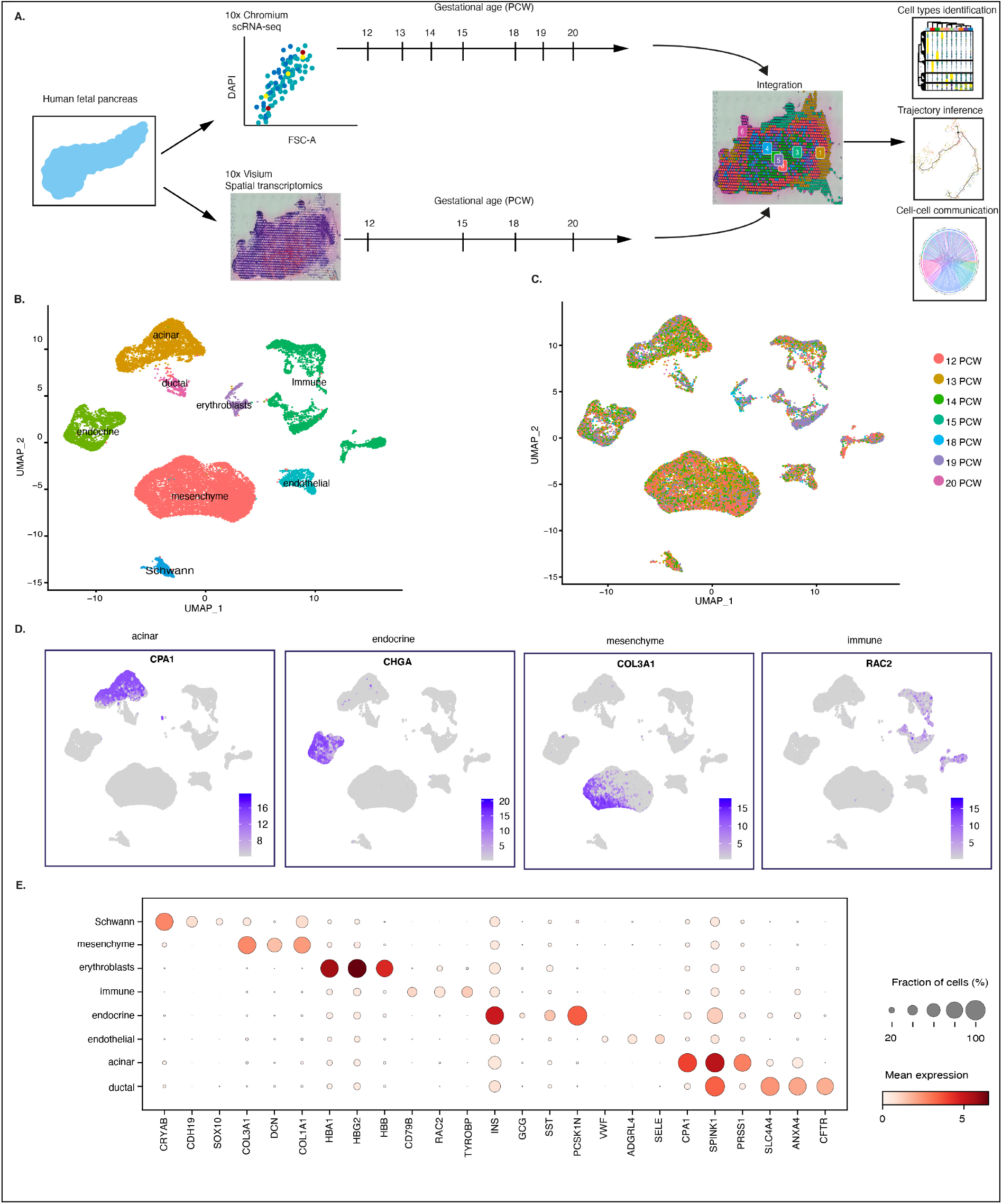
Single cell sequencing of the developing human pancreas. A) Schematic showing study design for spatiotemporal analysis of human pancreas development. B) UMAP embedding showing different annotated cell clusters in the developing human pancreas at 12-20 PCW. C) UMAP embedding of the developing pancreas transcriptomes showing the different gestational ages (PCW). D) Feature plots of selected marker genes for the major cell clusters. E) Dotplot showing marker gene distributions across the different cell populations.

### Spatial map of the developing human pancreas

It can be seen from Figure 1B and 1C that our scRNA-seq analysis revealed the presence of multiple pancreatic cell types within the developing human pancreas at the early stages of the second trimester. To spatially localise them and profile the gene expression dynamics of the cells in their morphological context, we performed 10x Visium spatial transcriptomics on 8 pancreas sections retrieved at 12, 15, 18 and 20 PCW using replicate tissue sections that were approximately 100μm apart from each other. We sequenced the samples to a median depth of 177.5 × 10^6^ reads (interquartile range 116.9-294.4 × 10^6^), which yielded a mean of 1692 genes per spot and 3395 unique molecular identifiers (UMIs) per spot (Suppl. Fig 2). Pre-processing and analysis of gene expression signatures in Seurat (Hafemeister and Satija, 2019) revealed 3-9 cell clusters (3 clusters at 12 PCW, 8 clusters at 15 PCW and 9 clusters each at 18 and 20 PCW), which correlated to distinct spatial locations within the tissues (Fig 2A). Annotation using marker genes indicated that some clusters contained multiple cells, as expected of the ∼55μm spatial resolution available when using 10x Visium. Using this approach we were able to stratify populations of endocrine cells (showing high expression of the islet hormones GHRL, SST and GCG), pancreatic/endocrine progenitors (expressing NKX6-1, SOX9 and HES1) (Seymour et al., 2007), endothelial cells (expressing VWF and ANGPT2), ductal/acinar (expressing CFTR and HES6), acinar (expressing HES6) and mesenchymal cells (expressing COL3A1 and VIM) (Fig 2B, 18 PCW samples). We then identified spatially proximal cells at 18PCW (Dries et al., 2021) and found that spatial neighbourhoods were shared by acinar and endocrine cells, by endocrine, ductal, acinar and pancreatic progenitors, and also by endothelial and mesenchymal cells, suggesting that these neighbouring cell populations are more likely to interact together (Fig 2C). We denoted interactions between cells of the same or different cell types as homo- or hetero-typic, respectively. Mesenchyme-mesenchyme pairs had the highest cell proximity score, indicating highly enriched cell-cell interactions, while endocrine cells and endothelial cells were predicted to preferentially undergo the highest number of homotypic interactions (Fig 2C). Spatial proximity profiling also indicated that the highest heterotypic interactions occurred between endocrine cells and pancreatic progenitors, with substantial interactions also predicted between ductal/acinar cells and pancreatic progenitors and also between acinar cells and endocrine cells (Fig 2C). We created spatial networks among the different cell types by connecting neighbouring cells through a Delaunay triangulation implemented in Giotto (Dries et al., 2021) and we defined a hub region as an area with the highest number of neighbouring cells expressing the same gene. This allowed us to identify spatially variable genes driving spatial trends across the pancreas, which correlated to distinct hub regions at 15PCW (Fig 2D and 2E) that became less prominent at 20PCW as the pancreas expands and the cells intermingle (Fig 2F and 2G). Thus, it can be seen at 15 PCW that there was co-localisation of expression of CLPS, which codes for the acinar protein colipase, and INS (Fig 2E), indicating that at this stage of development the exocrine and endocrine pancreas are not yet defined by the distinct anatomy that is observed post-natally. However, by 20 PCW the acinar and endocrine cells were spatially separated and discrete clusters of INS-expressing cells were evident, which most likely reflects the presence of islets scattered within the pancreas (Fig 2G). The majority of the spatially correlated genes that we identified are established canonical markers of the endocrine (INS, GCG, PCSK1N), mesenchyme (COL1A1, COL3A1) and acinar cell populations (CEL, CPA1) (Suppl. Fig 3A-3C), but other novel spatially correlated genes were also identified. For example, acyl-CoA thioesterase 7 (ACOT7) and ATP1A1 were spatially correlated with Schwann cell populations, DCN, ADAM33 and COX6A1 with the mesenchyme and ACTA2 with endothelial cells (Suppl. Fig 3D and 3E).

**Fig 2:**
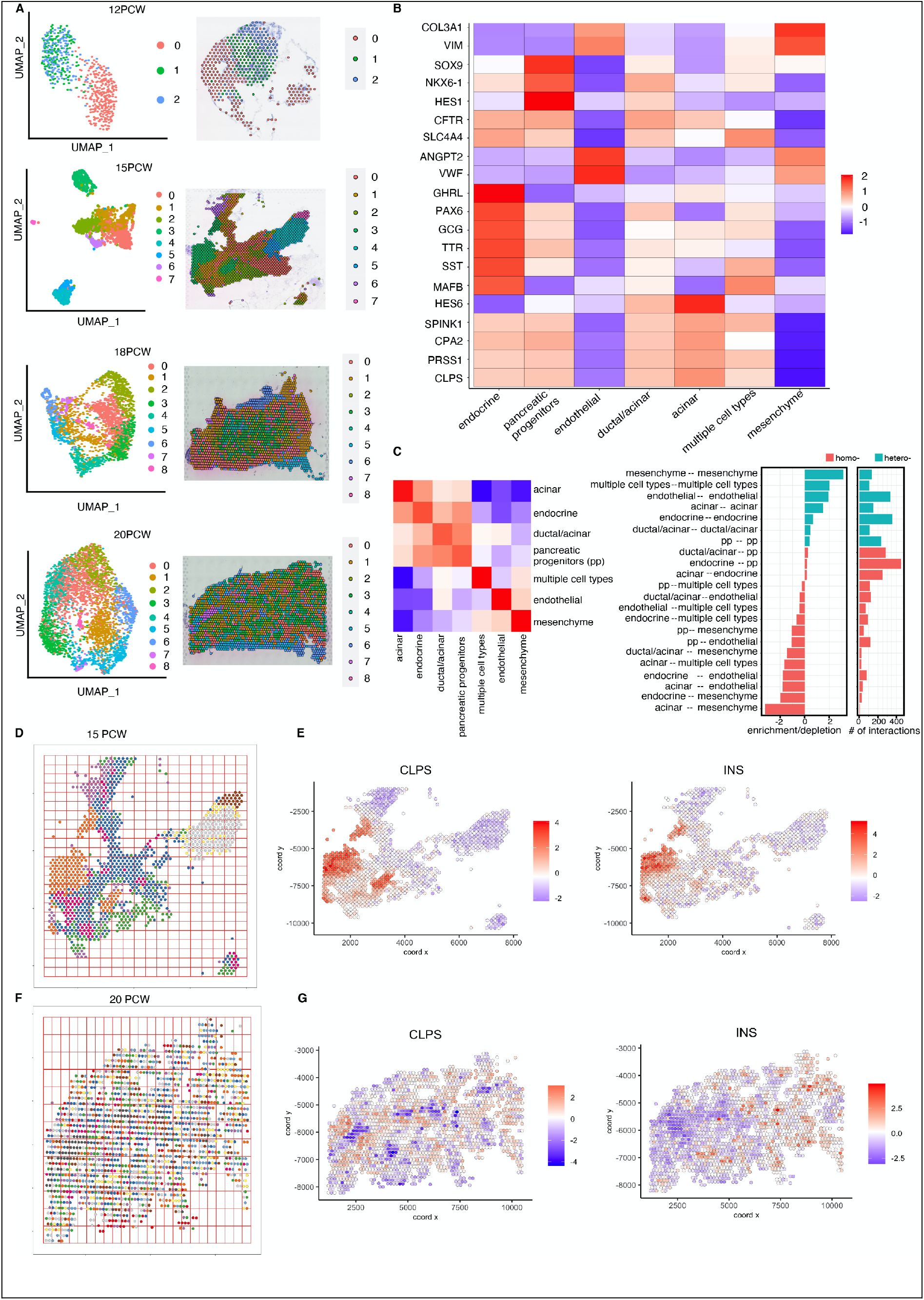
Spatial transcriptomics and cell-cell proximity map of the developing human pancreas. A) UMAP embedding and spatial projection of clusters on tissue slides at 12 PCW, 15PCW, 18PCW and 20PCW. B) Marker gene distribution and manual annotation of clusters at 18PCW. C) Heatmap (left) and bar chart (right) showing spatial proximity enrichment or depletion of cell type pairs and cell-cell interactions at 18PCW. D) Spatial grid containing cells based on their spatial locations at 15PCW. E) CLPS and INS gene expression at 15PCW showing exocrine and endocrine cells are not yet fully separated. F) Spatial grid containing cells based on their spatial locations at 20PCW. G) CLPS and INS gene expression indicating that exocrine and endocrine cells are spatially separated at 20PCW.

### Cell type deconvolution of the spatially resolved developing human pancreas transcriptome

While the spatial transcriptomics analysis provided novel information on cell-cell proximity in the human pancreas at different stages of development (Fig 2), the 55μm spot diameter of 10x Visium transcriptomics does not allow single cell resolution. We therefore combined our 10x Visium data with scRNA-seq data to characterise developing human pancreatic cell types at each spatial voxel, an approach that has recently been used to map the human endometrium (Garcia-Alonso et al., 2021) and the developing chicken heart (Mantri et al., 2021). We first integrated our scRNA-seq data (Fig 1) with a recently published dataset on the developing human pancreas at 8-19 WPC (Yu et al., 2021) using regularised negative binomial regression (Hafemeister and Satija, 2019) to increase the temporal resolution (Suppl. Fig 4A). To distinguish the two datasets, we refer to the integrated data as ‘combined scRNA-seq’. Following filtering and quality control, we retained 53,204 cells from 8 to 20 PCW. We found a strong correlation between the two datasets (r=0.8) (Suppl. Fig 4B) and combining the data allowed us to identify the clusters we had already found in our scRNA-seq dataset (Fig 1), but with improved resolution (Suppl. Fig 4C-4E). We deconvoluted the cell type composition at each spatial spot of the 10x Visium samples by projecting the combined scRNA-seq and spatial transcriptomics data into a common latent space and then we identified anchor cells that shared suffficient neighbourhood by canonical correlation analysis (Stuart et al., 2019). This allowed the transfer of cell annotations in the scRNA-seq data to the spatial transcriptomics data, thereby identifying alpha, beta, delta, endocrine progenitors, acinar, ductal, endothelial, Schwann, immune and mesenchymal cells *in situ* (Suppl. Fig 5). We then incorporated the cell type predictions into a deep-learning based method (Pham et al., 2020) to visualise the cell composition within each spatial voxel and predict cell proportions at each stage of development (Fig 3). As expected, we found an expansion in epithelial cells as the pancreas develops (e.g. a 175% increase in acinar cells at 20 PCW compared to 12 PCW) and a corresponding decrease in the proportion of mesenchymal cells (Fig 3A-D). Mesenchymal cells were mostly located at the pancreas periphery and most of the cell types existed in hub regions at 12 PCW, which was more prominent at 15 PCW (Fig 3A and 3B). For example, distinct regions of acinar, immune, mesenchyme, endocrine/endocrine progenitors and Schwann cells were evident at 15 PCW (Fig 3B), while by 20 PCW the cells were distributed throughout the pancreas, the bulk of which were acinar (Fig 3D). Among the endocrine cells, alpha cells were more than twice as abundant as beta cells at 12 PCW but at 15, 18 and 20 PCW there were comparable proportions of these two cell types. As adult human islets contain ∼55% beta cells and ∼38% alpha cells (Cabrera et al., 2006) it is likely that there is further expansion in beta cell numbers as gestation extends beyond 20 PCW. At all developmental stages that we interrogated, immune cells were spatially co-located with mesenchymal and endothelial cells but not with the endocrine cells (Fig 3A-D), while Schwann cells were in close spatial proximity to mesenchymal cells and endocrine progenitors at 15 PCW (Fig 3A and 3B).

**Fig 3:**
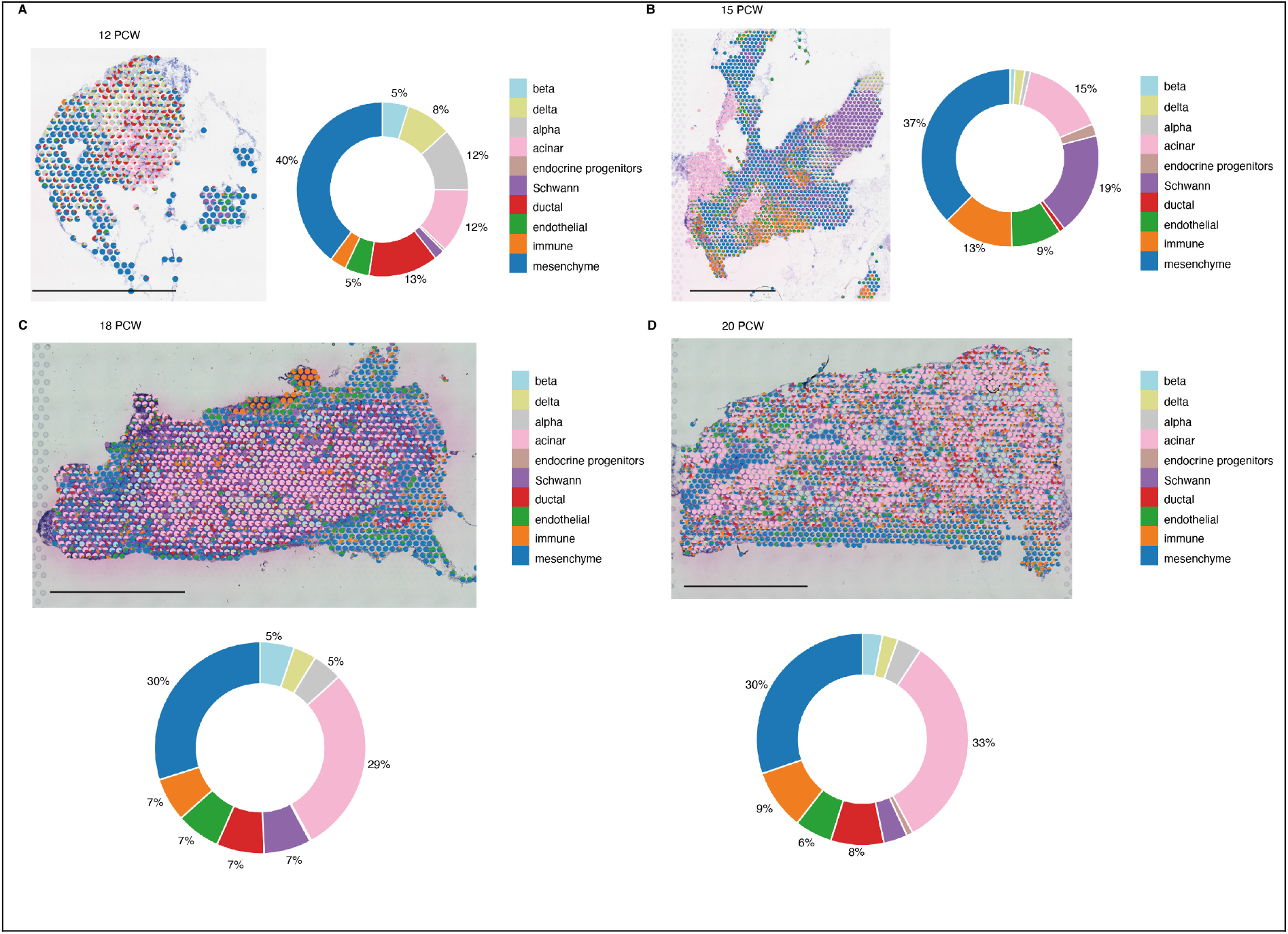
Cell-type map of the developing human pancreas. Cell type prediction overlay on ST spots and donut charts showing the proportion of cells inferred from scRNA-seq at A) 12 PCW, B) 15PCW, C) 18PCW and D) 20PCW. Scale bar at all stages = 2mm.

### Lineage dynamics within the endocrine compartment

Having identified the various cell types in our scRNA-seq and ST datasets, we next focused on gene expression trajectory analysis of the endocrine cells. Reclustering the endocrine component of our scRNA-seq data led to identification of 12 sub-clusters (Suppl. Fig 6A) and using differentially expressed genes and canonical markers we identified 3 endocrine progenitor populations (NEUROG3+), 4 beta cell populations (INS+), 2 delta cell clusters (SST+), and a cluster each for alpha (GCG+, which was also positive for PPY cells), and epsilon (GHRL+) cells (Suppl. Fig 6B and 6C). The three endocrine progenitor (EP) clusters were distinguished based on their gene expression: we denoted those with very high NEUROG3 expression as NEUROG3^hi^, those with low NEUROG3 and low insulin expression as NEUROG3^low^/INS^low^, suggestive of cells in transition towards beta cells, and those with very low expression of several islet hormones as polyhormonal (INS^low^/GCG^low^/SST^low^/GHRL^low^) (Fig 4A). We then sought to identify lineage relationships among the endocrine progenitor sub-clusters using time-series trajectory analysis (Tran and Bader, 2020), which showed that the three EP populations were found earlier in the inferred developmental timescale and were mostly from 12 and 13 PCW, as expected (Fig 4B). The trajectory analysis also predicted that cells in the NEUROG3^hi^ cluster were destined to transition to one of the beta cell clusters containing equal proportions of early- (12 and 13 PCW) and late-stage beta cells (18 PCW), while late-stage beta and delta cells (18-20PCW) and mid-stage alpha cells (14 and 15 PCW) were predicted to differentiate from NEUROG3^low^/INS^low^ EP populations (Fig 4B). The late-stage beta cells showed increasing expression of the maturity markers MAFA and UNC3 (Salinno et al., 2019; van der Meulen et al., 2012) (Suppl. Fig 6D). Some epsilon cells were clustered within an intermediate cluster along the NEUROG3^low^/INS^low^ EP trajectory as they transitioned to beta or delta cells (Fig 4B), possibly suggesting the multipotent nature of epsilon cells in development (Arnes et al., 2012). We then used Monocle 3 pseudotime analysis (Cao et al., 2019; Trapnell et al., 2014) to investigate whether we could recapitulate the trajectory predictions made using the time-series trajectory method. Monocle 3 identified an EP population with low NEUROG3 and low INS expression (Fig 4C), similar to NEUROG3^low^/INS^low^ EP (Fig 4B), and cells in this population also had low GCG expression. The cells also expressed important transcription factors known to be involved in endocrine pancreas development such as NEUROD1, insulin gene enhancer protein ISL1 and PAX6 (Fig 4D). In addition, genes that have been implicated in development of other tissues were also identified, such as insulin like growth factor binding protein like 1 (IGFPBL1), prothymosin alpha (PTMA) and G protein subunit gamma 8 (GNG8) (Emmanouilidou et al., 2013; Fujino et al., 2007; Gonda et al., 2007; Kriegebaum et al., 2010) (Fig 4D). The Monocle analysis indicated that the EP cells later split into two branches, the first of which was of beta cell lineage while the second branch contained both alpha and beta cells (Fig 4C). We investigated branch-dependent gene expression as the NEUROG3^low^/INS^low^ EP cells differentiated into alpha or beta cell lineages, and identified novel branch-specific genes in addition to established markers (Fig 4D), indicating that transcriptional maturation occurred over this developmental timeframe. To determine whether a similar endocrine progenitor trajectory exists in situ we identified cells that were positive for both NEUROG3 and INS at 20 PCW (Fig 4Ei) and ran spatial trajectory inference (Pham et al., 2020). This allowed us to explore progression towards endocrine cells (Fig 4Eii), and we identified genes that were upregulated or downregulated along the spatial trajectory (Fig 4Eiii). Known marker genes of endocrine cells such as chromogranin A and B (CHGA, CHGB), transthyretin (TTR) (Su et al., 2012), glucagon (GCG), somatostatin (SST), insulin (INS) and proprotein convertase subtilisin/kexin type 1 inhibitor (PCSK1N) were positively correlated with the spatial trajectory while the pancreatic progenitor surface marker glycoprotein 2 (GP2) (Cogger et al., 2017) and the acinar-related genes SPINK1, CLPS, CPA1 and PRSS1 were downregulated, indicating a transition of the progenitors to a more terminally differentiated endocrine state.

**Fig 4:**
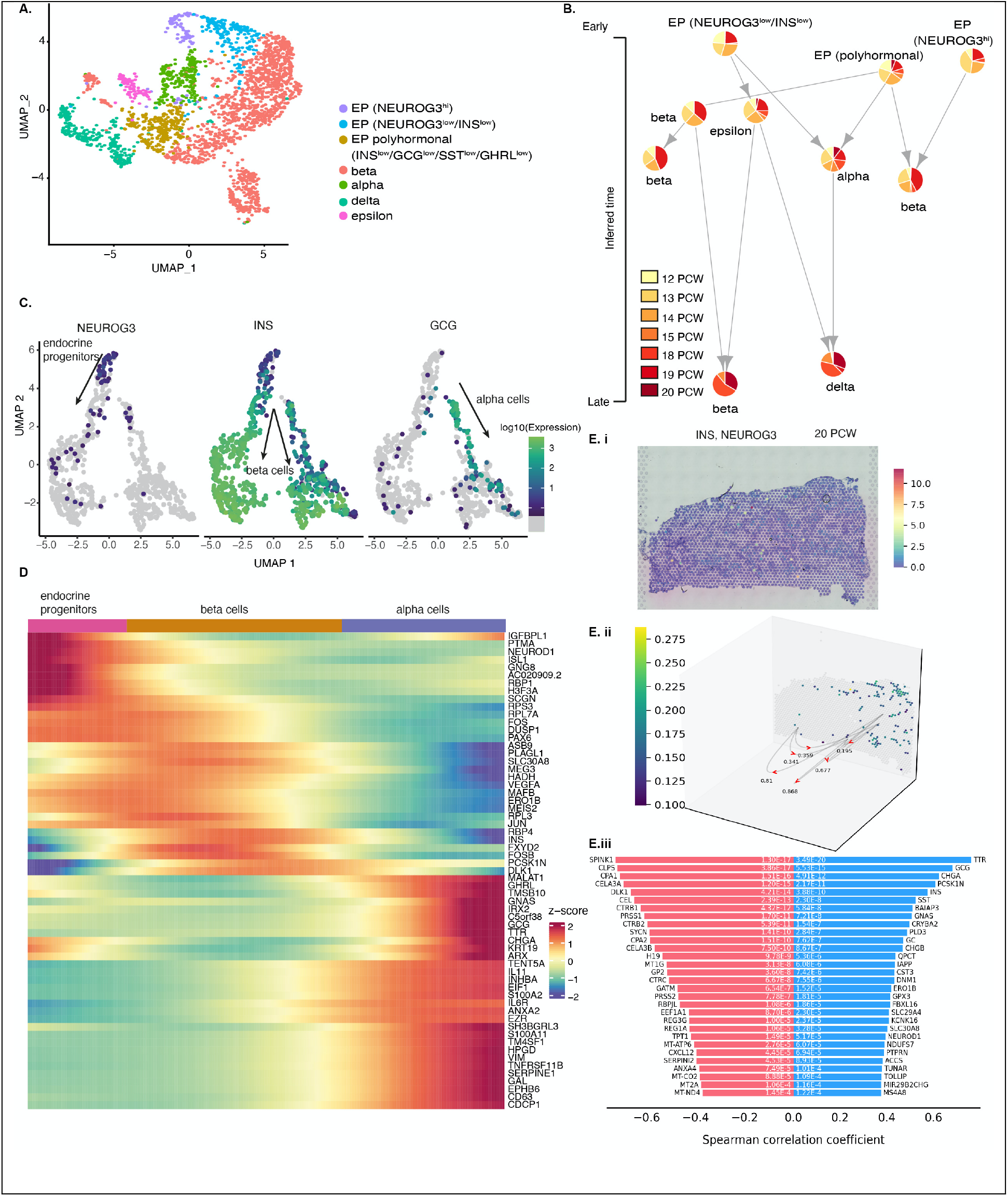
Identification of cell types and trajectory inference within the endocrine cluster. A) UMAP embedding showing annotated sub-clusters of the endocrine cells. B) Time-series trajectory inference of the EPs showing predicted transitions and lineage decisions into other endocrine cell types. C) Pseudotime analysis in Monocle 3 showing predicted bifurcation of NEUROG3^low^/INS^low^ endocrine progenitors into beta and alpha cell lineages. D) Heatmap showing branch-specific genes as endocrine progenitors differentiate towards beta and alpha cells. Ei) H&E image of 20 PCW pancreas showing cells positive for INS and NEUROG3. Eii) Pseudo-space-time analysis of an INS^+^/NEUROG3^+^ cluster showing trajectory directions Eiii) Top transition markers that are either positively correlated (blue) or negatively correlated with the spatial trajectory in Eii).

### Role of Schwann cells in endocrine cell development

After examining the lineage dynamics within the endocrine cluster, we next investigated the contribution of non-endocrine cells to endocrine specification. We focused on Schwann cells as they were spatially co-located with endocrine progenitor cells at 15 PCW (Fig 3B), suggesting that intercellular communications between these cells could contribute to endocrine cell differentiation. We identified five sub-clusters in our scRNA-seq data which all expressed the Schwann cell marker CRYAB (Suppl. Fig 7A and 7B). We annotated clusters 0, 1 and 4 as Schwann cell precursors (SCP) based on expression of the stem cell marker SOX2 (Liu et al., 2015) and cadherin 19 (CDH19) (Takahashi and Osumi, 2005), while cluster 3 was defined as an immature Schwann cell (iSC) population as it specifically expressed GAP43 (Jessen and Mirsky, 2005) (Suppl. Fig 7B). Cluster 2 expressed genes encoding myelin protein zero (MPZ) and proteolipid protein 1 (PLP1) (Suppl. Fig 7C) and gene ontology analysis of cluster 2 identified that these cells were associated with myelination and axon development (Suppl. Fig 7D) so they were designated myelinating Schwann cells (mSC).

Given the close proximity of Schwann cells with endocrine progenitors during human pancreas development (Fig 5A) we therefore investigated cell-cell connectivity (Raredon et al., 2021) in our scRNA-seq data to identify ligand-receptor pairs that may participate in paracrine signaling between these populations. We identified multiple ligand-receptor pairs between Schwann and endocrine progenitors (Fig 5B) and focused on the L1 cell adhesion molecule (L1CAM)-ephrin B2 (EPHB2) interaction, as L1CAM was the most highly predicted ligand and its expression was specific to Schwann cells where it was restricted to the iSC populations (Suppl. Fig 7C), while EPHB2 was expressed by endocrine progenitors and ductal cells (Suppl. Fig 7E). We mapped the expression of L1CAM-EPHB2 to our spatial transcriptomics data at 15 PCW (Pham et al., 2020) and identified that local hot spots of L1CAM-EPHB2 interactions occurred at the interface between Schwann cells and endocrine progenitors (Fig 5C). These observations are consistent with Schwann cells being spatially co-located with endocrine progenitors and contributing to beta cell maturation via the L1CAM-EPHB2 pathway.

**Fig 5:**
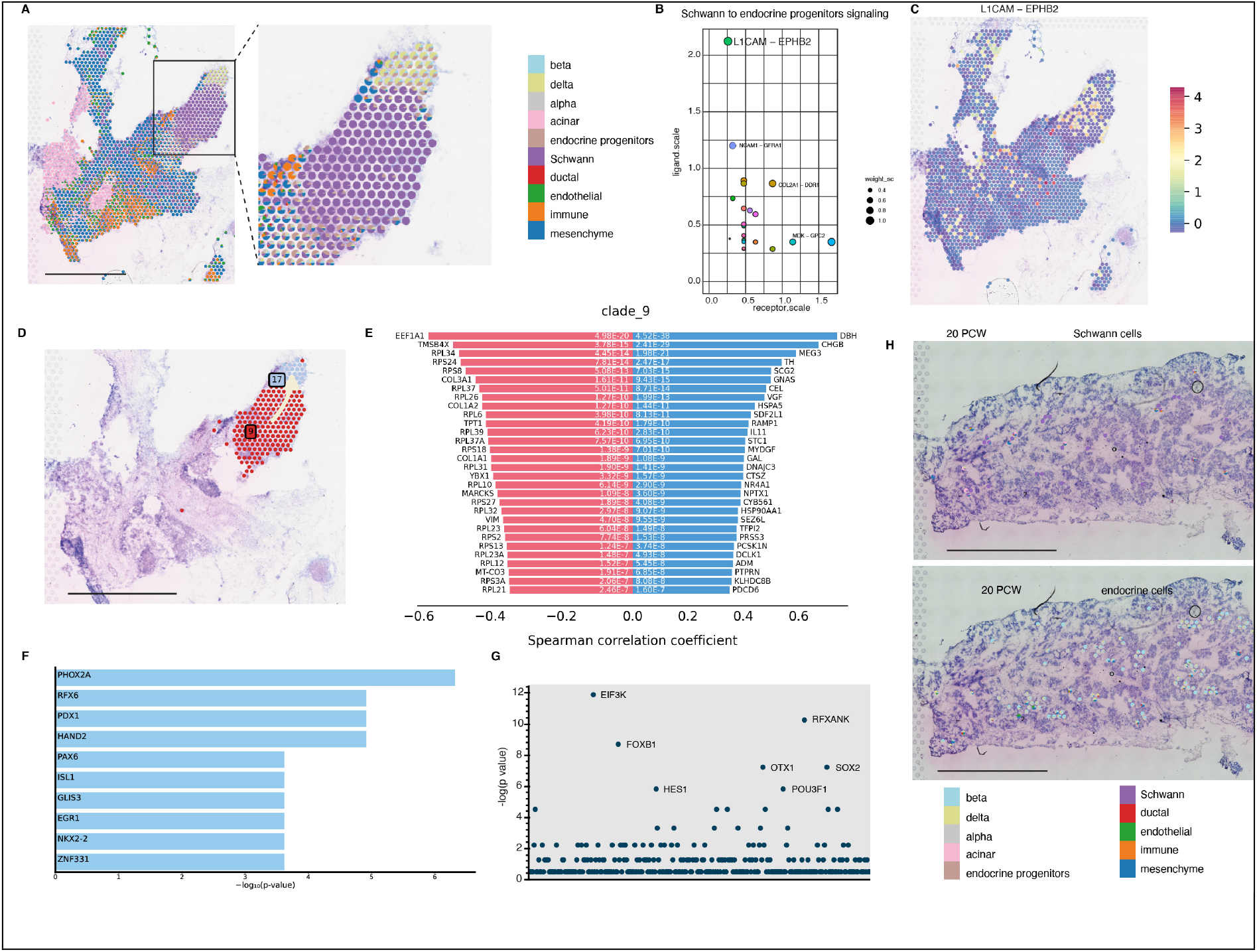
Role of Schwann cells in endocrine specification. A) scRNA-seq cell type prediction overlay on ST spots at 15 PCW. The zoomed in box shows spatial proximity of Schwann cells and endocrine cells. B) Ligand-receptor pairs predicted from scRNA-seq for Schwann-endocrine progenitor signalling. C) Spatial mapping of L1CAM-EPHB2 pairs in situ at 15 PCW. D) Visualisation of spatial trajectory between Schwann (clade 9) and endocrine clusters (clade 17). E) Top 30 transition markers that are positively correlated (blue) or negatively correlated (red) with the spatial trajectory between Schwann and endocrine clusters. F) Gene set enrichment analysis of transcription factors greater than three fold enriched in the positively correlated transition markers G) Manhattan plot showing -log10(p values) for transcription factor co-expression with the gene sets that were negatively correlated with the spatial trajectory. H) Visualisation of Schwann and endocrine cell clusters at 20 PCW indicating that the cells are no longer spatially co-localised.

We also used partition-based graph abstraction (PAGA) (Wolf et al., 2019), which shows unbiased lineage relationships based on gene expression among cell clusters, to investigate whether Schwann cells and endocrine progenitors shared lineage relationships. This analysis indicated that cluster 3 Schwann cells (Suppl. Fig 8A and 8B) shared connected edges with cluster 9 endocrine cells, containing endocrine progenitors, beta and delta cells (Suppl. Fig 8C) and these cell populations were close to each other within the reduced dimensionality space at 15 PCW (Suppl. Fig 8D). As expected, a diffusion pseudotime plot (Haghverdi et al., 2016) indicated that Schwann cells were transcriptionally earlier among all the clusters, so we set them as the root of the trajectory (Suppl. Fig 8E). We then performed spatial trajectory inference (Pham et al., 2020) on the Schwann cell cluster, and observed that endocrine cells were placed at the other end of Schwann cell trajectory (Fig 5D), with transition markers that were upregulated (blue) or downregulated (red) along the trajectory (Fig 5E). We performed gene set enrichment analysis (Kuleshov et al., 2016) on the upregulated transition markers and found significant enrichment (p<0.0001) of transcription factors involved in endocrine specification such as RFX6, PAX6, PDX1, ISL1 and NKX2-2 (Fig 5F). Transcription factors enriched in the gene sets that were downregulated or negatively correlated with the trajectory (i.e. genes whose expression was high in Schwann cells and low in endocrine cells) included SOX2, involved in stemness (Schaefer and Lengerke, 2020) and POU3F1, a neuronal fate gene (Zhu et al., 2014), suggesting that cells along the Schwann-endocrine trajectory are likely to lose their stemness and neuronal commitment. At 20 PCW, where endocrine cells were more terminally differentiated compared to 15 PCW, Schwann cells were no longer spatially co-localised with endocrine progenitors or other endocrine cells (Fig 5H).

### Mesenchymal heterogeneity in the developing human pancreas

Despite the important structural and biochemical roles that the mesenchyme plays in mouse pancreas organogenesis (Hibsher et al., 2016; Landsman et al., 2011), little is known about mesenchymal cell heterogeneity in the developing human pancreas. We therefore re-clustered the mesenchyme (10834 cells) and identified 17 transcriptionally distinct sub-clusters, all of which expressed the mesenchymal genes COL1A1, COL3A1 and VIM (Fig 6A & 6B). We annotated the clusters based on expression of known marker genes. Thus, clusters 0, 1, 2 and 4 were designated mesothelial cells based on expression of the Wilms’ tumour gene, WT1 (Ariza et al., 2018; Armstrong et al., 1993) and IGFBP2 (Namvar et al., 2018); clusters 12 and 15 were vascular smooth muscle (VSM) cells since they expressed alpha smooth muscle actin ACTA2 and transgelin TAGLN (Li et al., 1996; Majesky et al., 2011); cluster 13 was proliferative mesenchyme as defined by strong expression of STMN1 and MKi67; while cluster 14 belongs to the hematopoietic lineage based on expression of HBB. Cluster 4 was also positive for the osteogenic marker CLEC11A (Yue et al., 2016) and it is likely to have differentiated from mesothelial cells.

**Fig 6:**
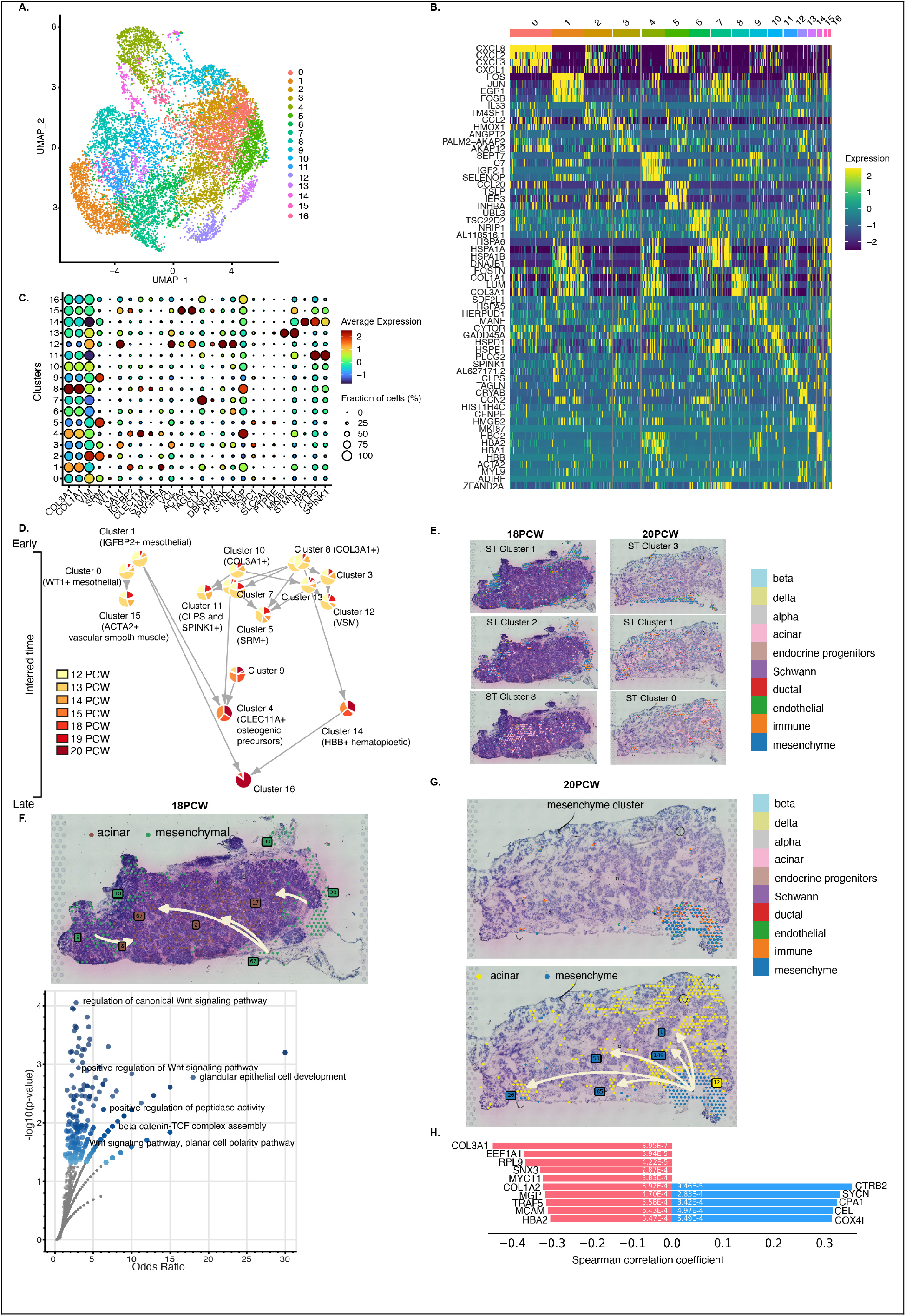
Heterogeneity and lineage predictions of the mesenchyme in the developing human pancreas. A) UMAP plot showing sub-clusters of the mesenchyme compartment. B) Heatmap showing differentially expressed genes in the mesenchyme 17 sub-clusters. C) Dotplot showing expression of marker genes across the clusters. D) Time-series trajectory inference showing lineage predictions within the mesenchyme. E) Visualisation of acinar and mesenchymal cell clusters at 18 and 20 PCW. F) Visualisation of mesenchymal-acinar trajectories with tissue localisation at 18 PCW. Pseudo-space-time analysis indicated three mesenchymal sub-clusters (clades 20, 66, 9) leading to four acinar sub-clusters (clades 17, 2, 67 and 8). The lower panel indicates a -log10(p-value) plot of gene ontology (GO) enrichment analysis on the positively correlated genes along the mesenchymal-acinar trajectories. GO pathways with p<0.05 are coloured blue. G) H&E image of a 20 PCW pancreas showing that mesenchymal and immune cells are spatially co-located within a cluster (upper panel). Visualisation of mesenchymal-acinar trajectories with tissue localisation at 20 PCW. Pseudo-space-time analysis indicated one mesenchymal cluster (clade 12) leading to multiple acinar sub-clusters (clades 1, 26, 50, 85 and 149) (lower panel). H) Top transition markers that are positively correlated (blue) or negatively correlated (red) with the mesenchymal-acinar spatial trajectories.

We identified that cluster 11 was enriched for the acinar genes colipase (CLPS) and serine protease inhibitor Kazal-type 1 (SPINK1) (Fig 6C). Given the co-expression of CLPS and SPINK1 within this mesenchyme cluster, we hypothesised an association between the mesenchyme and acinar maturation. Time-series trajectory analysis indicated that IGFBP2-expressing cluster 1 mesothelial cells represented an early cell state in the lineage, consisting mainly of cells at 12 and 13 PCW, and these cells transitioned to become cluster 15 VSM cells that expressed ACTA2 (Fig 6D). This is consistent with mesothelial cells being well-established precursors for VSM (Que et al., 2008; Wilm et al., 2005) and they have also been proposed to contribute to the VSM lineage in mouse pancreas (Byrnes et al., 2018). Clusters 8 and 10 strongly expressed the pan-mesenchymal marker COL3A1 and they were predicted to differentiate into multiple lineages including cluster 12 VSM, cluster 14 hematopioetic cells, cluster 4 osteogenic precursors and spermidine-expressing cluster 5 cells. The analysis placed the CLPS- and SPINK1-expressing cluster at the exit of a trajectory (Fig 6D), suggesting a likely lineage relationship between the mesenchyme and acinar cell populations. We therefore investigated lineage dynamics between mesenchymal and acinar cells in situ using our spatial transcriptomics data. This indicated that at 18 and 20 PCW mesenchymal cells were mostly located at the periphery of the pancreas and were often in association with endothelial or immune cells, while acinar cells were adjacent to them (Fig 6E). Spatial trajectory analysis predicted a transition from mesenchymal to acinar cells at 18 PCW (Fig 6F, upper panel) and at 20 PCW (Fig 6G). We identified transition genes that were upregulated along the spatial trajectory from mesenchymal clades 9, 20 and 66 to acinar clades 8, 17, 2 and 67 respectively (Fig 6F, upper panel, Table 6). Enrichment analysis (Kuleshov et al., 2016) of the upregulated transition genes showed significant enrichment of genes involved in Wnt/beta-catenin signalling, planar cell polarity and glandular epithelial cell development (Fig 6F, lower panel), indicative of processes involved in acinar cell development and growth (Murtaugh, 2008; Wells et al., 2007). Our trajectory-based differential gene expression analysis at 20 PCW also identified that mesenchymal genes including COL3A1, COL1A2 and MGP were downregulated while expression of the acinar genes CPA1 and CEL was increased (Fig 6H).

## Discussion

Co-ordinated interactions between different cells at critical time windows is essential for organogenesis, yet our understanding of how the diverse human pancreatic cell types interact during development is limited. Using scRNA-seq and spatial transcriptomics, we identified and localised endocrine, acinar, ductal, endothelial, immune, Schwann and mesenchymal cell types in human fetal pancreas at 12-20 gestation weeks. To improve the temporal resolution of our scRNA-seq, we integrated it with a recently published dataset on the developing human pancreas (Yu et al., 2021) and used the combined data to deconvolute the cell types in our 10x Visium samples. Our data revealed varying proportions of pancreatic cell types, with mesenchymal cells decreasing in number over developmental time coupled to a corresponding increase in epithelial cell populations. We uncovered heterogeneity within the endocrine, Schwann and mesenchyme compartments with predicted lineage relationships among the sub-populations. Major strengths of our study are the availability of fetal pancreases over the relatively wide developmental range of 12-20 PCW and our combination of scRNA-seq and spatial transcriptomics over this time-course. This has enabled us, for the first time, to reconstruct developmental trajectories occurring in situ, and to delineate the contribution of the mesenchyme and Schwann cells in the differentiation towards acinar and endocrine lineages, respectively.

It has been suggested that neural crest cells, the precursors of Schwann cells, directly differentiate into islet cells in mice (Pearse and Polak, 1971), but rats that lack Schwann cells progenitors do not fail to develop an endocrine pancreas (Pictet et al., 1976). Nonetheless, the plasticity of Schwann cell precursors in pancreas development is not fully appreciated and they have been implicated in cell specification in other tissues (Perera and Kerosuo, 2021). We have shown here that a Schwann cell subset, the immature Schwann cells, express L1CAM and are located in spatial proximity to endocrine progenitors, with local hotspots of L1CAM-EPHB2 interactions. We therefore propose that the pancreatic immature Schwann cells contribute to maturation of endocrine progenitor cells via L1CAM-EPHB2 signalling, and this may be important in improving beta cell mass since neural crest cell/beta cell co-transplantation has been shown to increase beta cell proliferation and improve normoglycaemia in diabetic mice (Olerud et al., 2009).

We have also uncovered underappreciated heterogeneity within the mesenchyme, which is reminiscent of the diversity of mouse pancreas mesenchyme (Byrnes et al., 2018). Notably, we identified an acinar cluster within the mesenchyme that was placed at the end of the trajectory of a pan-mesenchymal COL3A1+ cluster. A similar mesenchyme-acinar trajectory was predicted in situ from our spatial transcriptomics data, and we identified significant enrichment of gene ontologies involved in acinar cell differentiation, such as positive regulation of canonical Wnt-beta-catenin signalling, planar cell polarity and glandular epithelial cell development (Murtaugh, 2008; Wells et al., 2007). This points to the importance of the mesenchyme in appropriate differentiation of acinar cells in the developing human pancreas, which is in agreement with a previous study where mouse exocrine pancreas failed to develop in the absence of pancreatic mesenchyme (Gittes et al., 1996).

In summary, we have characterised and spatially resolved multiple human pancreatic cell populations at multiple developmental stages. We have identified sub-populations of human endocrine progenitors, novel genes that may direct their differentiation to beta or alpha cell lineage, and the influence of pancreas microenvironment on endocrine progenitor differentiation. Our data also identified the roles of Schwann precursor cells and mesenchymal cells in the differentiation of endocrine progenitors and acinar cells, respectively. We have also provided an interactive web resource (www.humanpancreasdevelopment.org) to explore the multi-dimensional data presented in this study. Further exploration of the endocrine lineage relationships and functional roles of the branch-dependent genes along the beta cell lineage will be important in optimising protocols to generate functional beta cells in vitro.

## MATERIALS AND METHODS

### RESOURCE AVAILABILITY

#### Lead contact

Further information and request for reagents should be directed to and will be fulfilled by the lead contact, Professor Shanta Persaud (shanta.persaud{@}kcl.ac.uk). We did not generate any unique reagents from this study.

#### Data and code availability

The raw data used for this study will be deposited on GEO and accession numbers will be provided. Single-cell RNA sequencing and spatial transcriptomics data were analysed using publicly available software. All R and python scripts will be uploaded at https://github.com/olaniru/human-fetal-pancreas.

### EXPERIMENTAL MODELS AND SUBJECT DETAILS

#### Human samples

Human embryonic and fetal pancreases were obtained from the MRC-Wellcome Trust Human Development Biology Resource, London, with appropriate ethical approval. Samples were transferred immediately after provision in cold Lebowitz medium (L-15) to KCL and processed immediately.

### METHOD DETAILS

#### Human fetal pancreas scRNA-seq dissociation protocol

On the day of retrieval, pancreases were rinsed in ice-cold PBS and any extra-pancreatic tissues were dissected out before they were inflated with 1mg/ml collagenase from *Clostridium histolyticum* (C2674). The tissues were digested at 37°C for 5 min (12-14 PCW) or 10 min (15-20 PCW), gently disrupted by pipetting with a wide-bore Rainin pipette and incubated in trypsin/EDTA for an additional 5 min at 37°C. Enzyme action was stopped with 10% FBS in RPMI medium and the samples were filtered through a 30μm mesh before being centrifuged at 1100rpm for 5min. Cell pellets were resuspended in RPMI containing 10% FBS. Cell number and viability were determined using a Countess II automated cell counter. At least 400,000 cells per pancreas with viability of 94±2% were carried forward for hash-tag staining. We initially investigated whether hash-tag (HTO) conjugation is feasible in the developing human pancreas by staining for the ubiquitous surface markers β2M and CD298 to which the oligonucleotide-tagged antibodies bind (Stoeckius et al., 2018). ∼450,000-2,000,000 cells were incubated for 10min in human TruStain FcX (10%v/v) to block non-specific binding before being exposed to varying concentrations (0.25, 0.5, 1.0 and 2μg/μl) of PE β2M/PE CD298 (BioLegend, mixed 1:1) for 30 min in the dark at 4°C, then washed with cold PBS containing 0.04% BSA. 0.1ng/μl DAPI was added and FACs sorting on a BD FACSAria 3 was used to determine the percentage of live β2M/CD298-positive cells. Subsequently, HTO staining was carried out with 0.5μg of β2M and CD298 antibody-conjugated oligonucleotides (Total-seq B Hastags 4, 6 and 8; BioLegend) with ∼500,000-750,000 isolated single cells. For each scRNA-seq run, about 60,000 live cells were sorted into RPMI containing 5% FBS (4°C) before proceeding to 10x Chromium library preparation and sequencing.

#### 10x Chromium scRNA-seq library preparation and raw sequence data processing

We used the 10x Genomics single cell RNA sequencing 3’ v3 chemistry for generating our scRNA-seq data. Isolated single cells at multiple gestational stages were stained individually with Totalseq B HTOs, pooled together into 4 batches (12, 13 and 13 PCW; 12, 14 and 19 PCW; 13, 14 and 19 PCW; 15, 18 and 20 PCW) and 20,000 cells from each batch were loaded on the 10x chip. Single cell mRNA and antibody (HTO) libraries were prepared according to the manufacturer’s protocol and paired-end sequencing was carried out on an Illumina NextSeq 2000, except for batch 15, 18 and 20 PCW which was sequenced on a HiSeq 2500. Generation of FASTQ files and alignment of raw sequencing reads with GRCh38 human genome reference data were carried out using CellRanger (version 6.0.0, 10x Genomics).

#### Hashed sample de-multiplexing

We first filtered the HTO antibody UMI count matrices to retain only those containing 10x cellular barcodes. We then used the HTODemux function in Seurat to separate the pooled samples into their individual components, as previously described (Stoeckius et al., 2018). We obtained a subset of the UMI count matrix and HTO count matrix using cellular barcodes contained in both matrices, which were log normalised and clustered using the clara k-mediod function. A negative binomial distribution was then fit to each cluster, with a default positive threshold of 99^th^ percentile. This threshold was used to determine whether the cells were positive or negative for the hashtag: cells positive for more than one hashtags were classed as doublets and were removed from analysis. We also performed genetic demultiplexing (Huang et al., 2019) and observed nearly 100% concordance with the de-hashing algorithm in Seurat.

#### 10x scRNA-seq data analysis

Output from Cellranger for the four scRNA-seq batches were loaded into Seurat (version 4.0.2) and merged. After filtering out low quality reads using nUMI > 500, nGene > 250, log10GenesPerUMI > 0.75 and mitoRatio < 0.25 thresholds, the Seurat objects were normalised individually by regularised negative binomial regression (SCTransform) to correct for batch effect and other technical variabilities (Hafemeister and Satija, 2019). The data were integrated using the IntegrateData function after selecting the most variable features which were used to extract integrating anchors using FindIntegrationAnchors. We then performed principal component analysis and used the first 20 dimensions to compute a Uniform Manifold Approximation and Projection (UMAP) (Becht et al., 2018). By FindNeighbors and FindClusters commands in Seurat, cells were clustered in a graph-based approach by the Louvain algorithm. Wilcoxon rank sum tests were performed to identify differentially expressed genes in each cluster using the FindAllMarkers function in Seurat. The clusters were annotated based on marker gene expression as endocrine (INS, GCG, SST, PAX6), acinar (CPA1, PRSS1, CLPS), ductal (SLC4A4, CFTR, ANXA4), mesenchymal (COL3A1, DCN, VIM), immune (RAC2, LYZ, TRAC), endothelial (VWF, ADGRL4, ANGPT2), Schwann (CRYAB, CDH19, SOX10) and erythroblasts (HBB, HBG2, HBA1).

#### Subclustering of clusters of interest

Endocrine, mesenchymal and Schwann clusters were first individually isolated using the Subset function in Seurat and re-analysed as described above (10X sc-RNA-seq data analysis). Endocrine subclusters were annotated with marker genes INS, GCG, SST, PYY, GHRL and NEUROG3, mesenchymal subclusters were annotated with COL3A1, COL1A1, WT1, IGFBP2, ACTA2, TAGLN, HBB, CLEC11A and MKi67 while Schwann subclusters were identified as Schwann cell precursors (SOX2 and CDH19), immature Schwann cells (GAP43) and myelinating Schwann cells (MPZ, PLP1).

#### Integration of scRNA-seq data with previously published 10x Chromium data

The scRNA-seq generated in this study were integrated with 10x Chromium data from a previous study on human fetal pancreas development (Yu et al., 2021), which was accessed from (https://ngdc.cncb.ac.cn/omix) with accession number OMIX236. To allow direct comparison with our data we only extracted data for the whole pancreas and excluded datasets that had been enriched for epithelial cells. We preprocessed each dataset by SCTransform and used the most variable features to identify integration anchors, which were then passed to the IntegrateData function to return a Seurat object containing an integrated expression matrix for all cells. The data were jointly analysed and visualised by UMAP. The clusters were annotated using marker genes as described above (10X sc-RNA-seq data analysis).

#### Spatial transcriptomics and raw sequence data processing

Spatial transcriptomics was carried out using the 10x Genomics Visium platform. 10μm tissue sections from OCT-embedded fresh frozen human pancreases at 12, 15, 18 and 20 PCW were mounted onto Visium Spatial slides and the sections were permeabilised for 30min to release mRNAs, which bind to the spatially barcoded-oligos present in the underlying spots and reverse transcribed, according to the manufacturer’s protocol. Libraries prepared from the cDNAs were sequenced on the Illumina NextSeq 2000 platform at >50,000 reads per spot generating >400M reads per section. Spaceranger software (version 3.1.0, 10x Genomics) was used to align and obtain raw counts from each of the spots on the Visium spatial transcriptomics slides against the GRCh38 human genome reference data.

#### 10x Visium spatial transcriptomics data analysis

The spatial transcriptomics raw gene expression matrix, together with spatial location of spots and tissue H&E images, were used to create a Seurat object with a Load10X_spatial function. After normalisation by SCTransform, we performed principal component analysis and reduced the dimensions to the top 20 principal components. Marker gene detection and differential gene expression were carried out using the FindAllMarkers function in Seurat. Genes that varied with locations in situ were identified using the FindSpatiallyVariableFeatures function, using default settings.

#### Spatial transcriptomics deconvolution and visualisation

To spatially map the different developing human pancreas cells in situ, the combined scRNA-seq datasets were integrated with 10x Visium spatial transciptomics data using the anchor-based integration pipeline in Seurat, which allowed the transfer of cell-type annotations from scRNA-seq to spatial trancriptomics. The cell type predictions from Seurat were loaded into a python package stLearn (Pham et al., 2020), where the cell types in every spatial spot were annotated and visualised as donut charts.

#### Spatial co-localisation of receptor-ligand pairs

To investigate cell-cell interactions between endocrine progenitors and Schwann cells, we first annotated cell type diversity within each spot using cell type predictions in Seurat. We then identified significant ligand-receptor pairs between neighbouring spots using CellPhoneDB (Efremova et al., 2020), as implemented in stLearn. Ligand-receptor hotspots were defined as spatial regions containing high numbers of interacting cells and high ligand-receptor co-expression.

#### Spatial transcriptomics analysis using the Giotto package

To create a spatial network and identify spatial co-expression patterns, we reanalysed the raw spatial gene expression matrix with Giotto (Dries et al., 2021), and retained only spots that overlapped with the tissue area. We removed genes of low expression and low quality spots using filterGiotto with default parameters. Following normalisation, we identified highly variable genes which were used to perform PCA analysis and shared-nearest neighbour identities were computed using the first 10 principal components. We performed Leiden clustering at a resolution of 0.4 to identify clusters of spots. We identified spatial co-expression modules and created spatial networks using the functions detectSpatialCorGenes and createSpatialNetwork, respectively.

#### scRNA-seq cell-cell communication analysis

To infer cell-cell communication and identify ligands and receptors involved in endocrine progenitor/ Schwann cell interactions, we used the Connectome v1.0.1 package (Raredon et al., 2021). Using the CellCellScatter function, we identified the top signalling ligand-receptor pairs between endocrine progenitors and Schwann cells.

#### Monocle 3 pseudotime trajectory analysis

Trajectory inference with Monocle 3 (Cao et al., 2019) was carried out on the endocrine Seurat subset. We set NEUROG3-positive cells as the root of the trajectory and identified differentially expressed genes across the trajectories using the Moran’s I test as implemented in Monocle 3.

#### Time-series trajectory analysis

Using gestational stages as time input, we used Tempora (Tran and Bader, 2020) to identify cell type relationships at different stages of pancreas development. Already processed Seurat objects were loaded and pathway enrichment profiles of the clusters were calculated using gene set variation analysis (Hänzelmann et al., 2013). Using the top 6 principal components, trajectories were built based on the clusters’ pathway enrichment profiles and visualised as piecharts showing the proportion of cells at multiple time points and arrows connecting each piechart were used to show lineage relationships.

#### Gene ontology enrichment analysis

Gene lists generated from the above-described analyses were used as inputs for gene ontology enrichment analysis, which was carried out on a web-based platform, Enrichr (Kuleshov et al., 2016).

#### Spatial trajectory analysis

To investigate cellular trajectories in situ, we reprocessed the raw spatial data following documented protocols in stLearn (Pham et al., 2020). Following Louvain clustering, we performed global and local pseudo-space-time trajectory analysis in stLearn which incorporates PAGA (Wolf et al., 2019) and a diffusion pseudotime method (Haghverdi et al., 2016) to reconstrust trajectories using changes in transcriptional states between clusters of interest. Genes that were differentially upregulated or downregulated along the trajectories were determined by Spearman’s rank correlation with a threshold of 0.3.

## Acknowledgements

We thank the Joint MRC/Wellcome Trust (grant# MR/R006237/1) Human Developmental Biology Resource (http://hdbr.org) for provision of the human fetal pancreatic material.

We acknowledge assistance from NIHR Guy’s and St Thomas’ Biomedical Research Centre Flow Cytometry Core Facility and Genomics Centre. O.E.O. was supported by a Novo Nordisk UK Research Foundation Grant and an NC3Rs Early Career Support Award.

## Conflicts of interest

The authors declare that they have no conflict of interest.

## Author contributions

Research design: OEO, SJP. Conducted experiments: OEO. Prepared scRNA-seq and 10x Visium libraries: HV, KRF, PD. Performed data analysis: OEO, UK, SK, SJP. Wrote or contributed to the writing of the manuscript: OEO, SJP. All authors revised the manuscript critically for important intellectual content and gave their approval for the current version to be published. OEO and SJP are the guarantors of this work and, as such, had full access to all the data in the study and take responsibility for the integrity of the data and the accuracy of the data analysis.

## Supplementary Figures

**Supplementary Figure 1:**
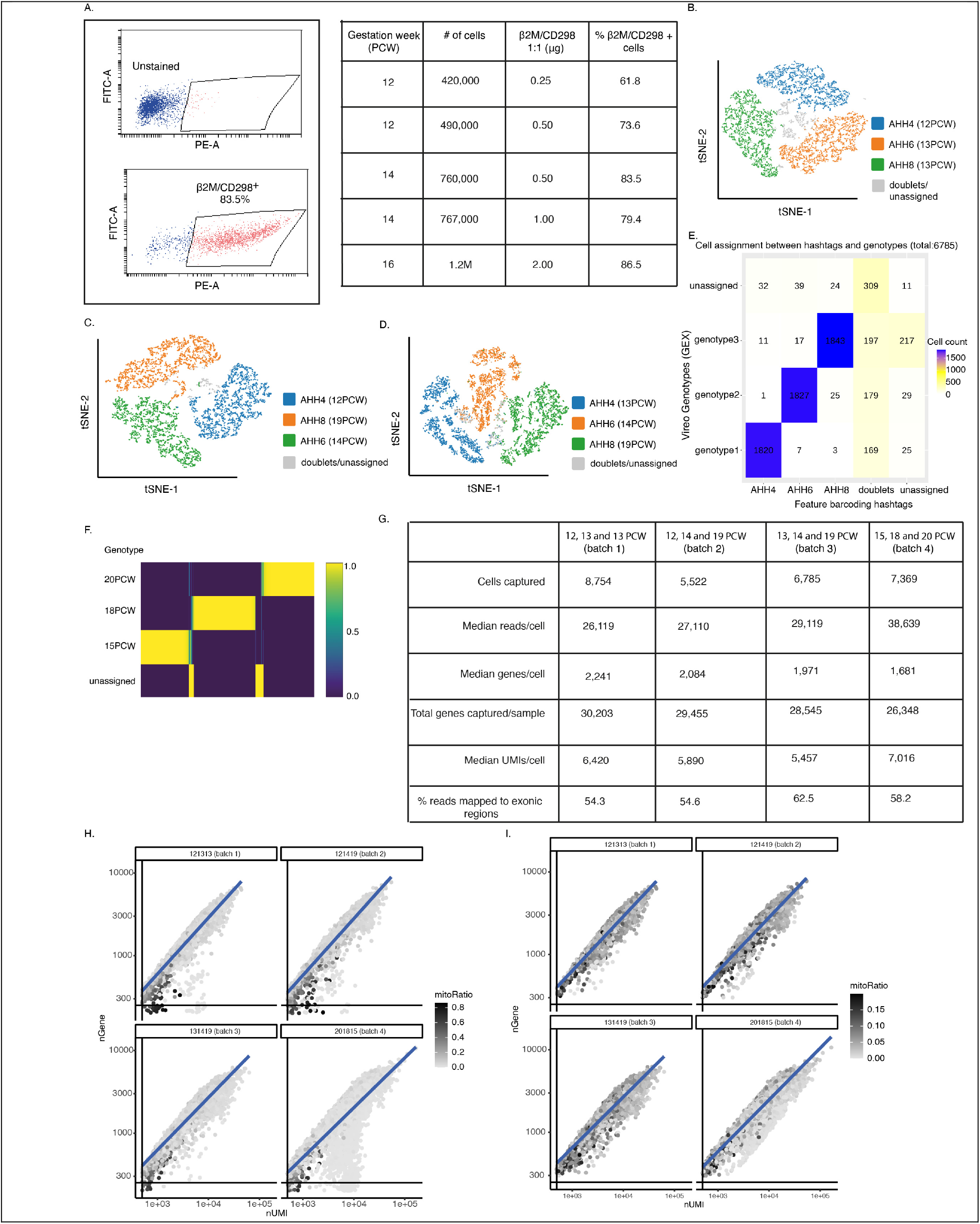
Demultiplexing and quality control of single cell RNA sequencing batches. **A)** Flow cytometry analysis of CD298 and β2M expression in human fetal pancreas at 14 PCW. The table shows the percentage of CD298+ and β2M+ cells at different gestation stages (12-16 PCW). **B)** t-distributed stochastic neighbourhood embedding (tSNE) of cells from hashtag (HTO) demultiplexing of a pool of single cells from 12, 13 and 13 PCW into their individual components. AHH4, AHH6 and AHH8 are the Total-seq B Hastags 4, 6 and 8 respectively. **C)** tSNE visualisation of cells from a 12,14 and 19 PCW pool using the HTO de-hashing algorithm. **D)** tSNE visualisation of cells from a 13,14 and 19 PCW pool using the HTO de-hashing algorithm. **E)** Genotype demultiplexing by Vireo (Huang et al., 2019) showed 98% concordance with HTO re-assignments of the single cells from the same pool of cells from 13, 14 and 19 PCW samples. The de-hashing method predicted a doublet rate of 12% while the number of unassigned cells (doublets and negatives) went down to 6.1% (415 cells) with genotye demultiplexing. **F)** Genotype demultiplexing of single cells from a pool of 15, 18 and 20 PCW cells into their individual components. **G)** Quality control metrics for the four batches of scRNA-seq performed in this study using 10x Chromium 3’ v3. **H)** Pre-filtering visualisation of the correlation between number of genes detected and the number of UMI per cell coloured by the fraction of mitochondrial reads. **I)** Post-filtering visualisation of the correlation between number of genes detected and the number of UMI per cell coloured by the fraction of mitochondrial reads. Low quality reads were filtered out using the following thresholds: nUMI > 500, nGene > 250, log10GenesPerUMI > 0.75 and mitoRatio < 0.25.

**Supplementary Figure 2:**
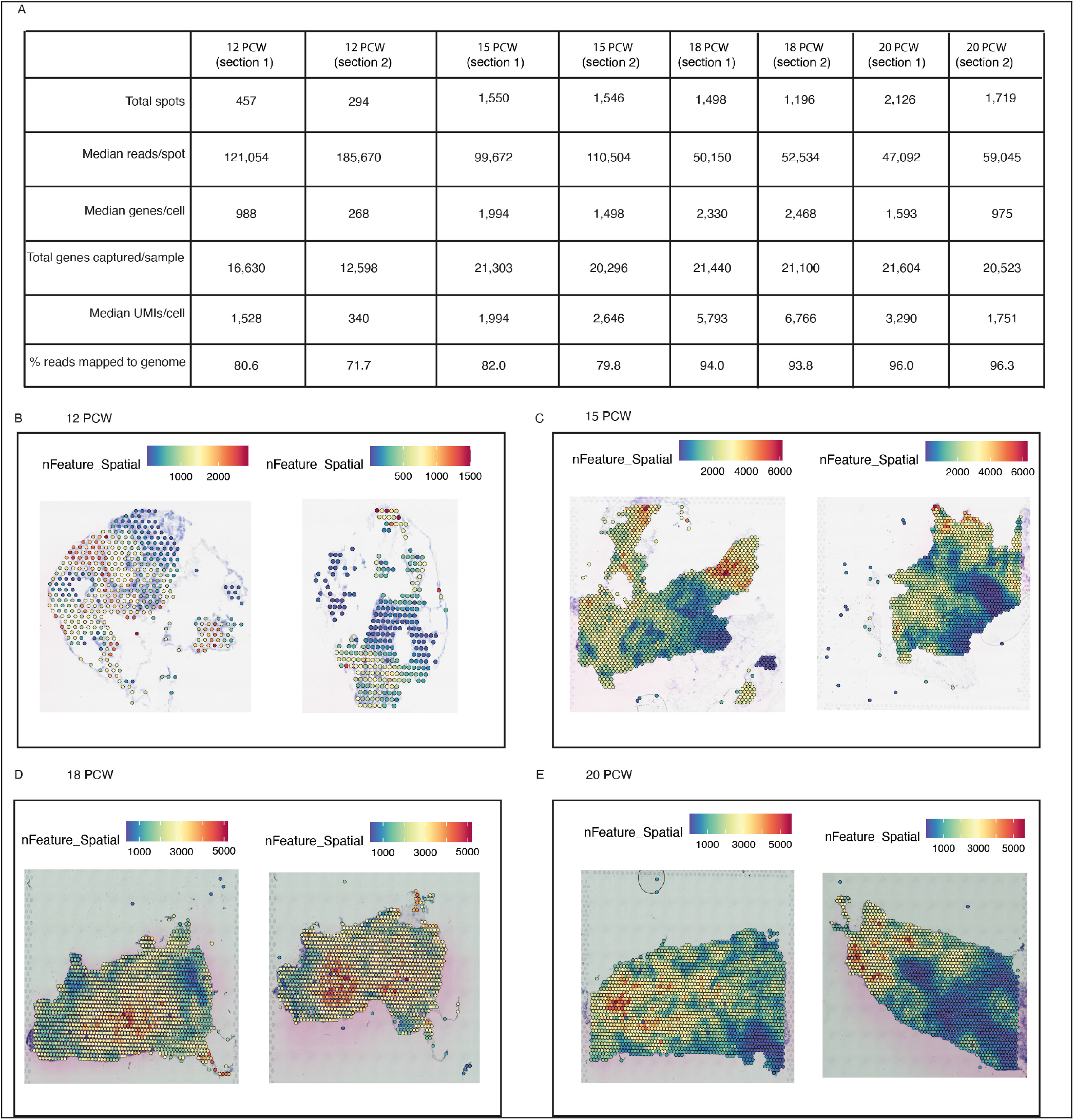
Summary of 10x Visium spatial transcriptomic data. **A)** Quality control metrics for the eight pancreas sections processed using 10x Visium. **B)** Spatial gene expression distribution across two pancreas sections (left and right) at 12 PCW, and at **C)** 15 PCW **D)** 18 PCW **E)** 20 PCW

**Supplementary Figure 3:**
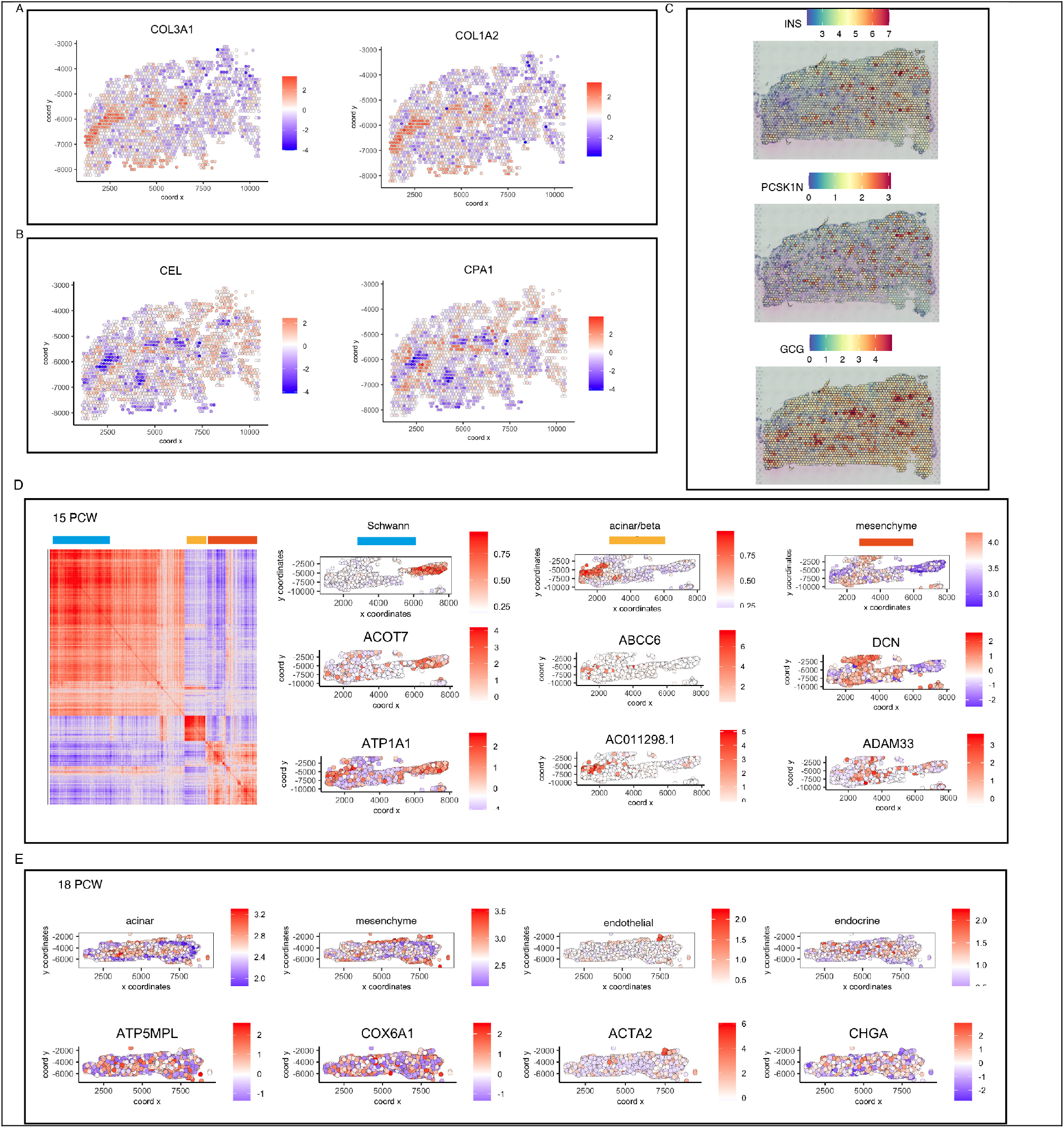
Spatially correlated genes in human pancreas development. **A)** Examples of mesenchyme canonical markers which were also identified as spatially variable genes. **B)** Examples of acinar canonical markers which were also identified as spatially variable genes. **C)** Examples of endocrine markers which were also identified as spatially variable genes. **D)** Heatmap showing spatial gene co-expression in the developing pancreas at 15 PCW. Three of the identified spatial co-expression modules are shown with different colours on top of the heatmap. To the right of the heatmap are examples of genes within the modules. **E)** Examples of identified spatial genes within the acinar, mesenchymal, endothelial and endocrine regions at 18 PCW.

**Supplementary Figure 4.**
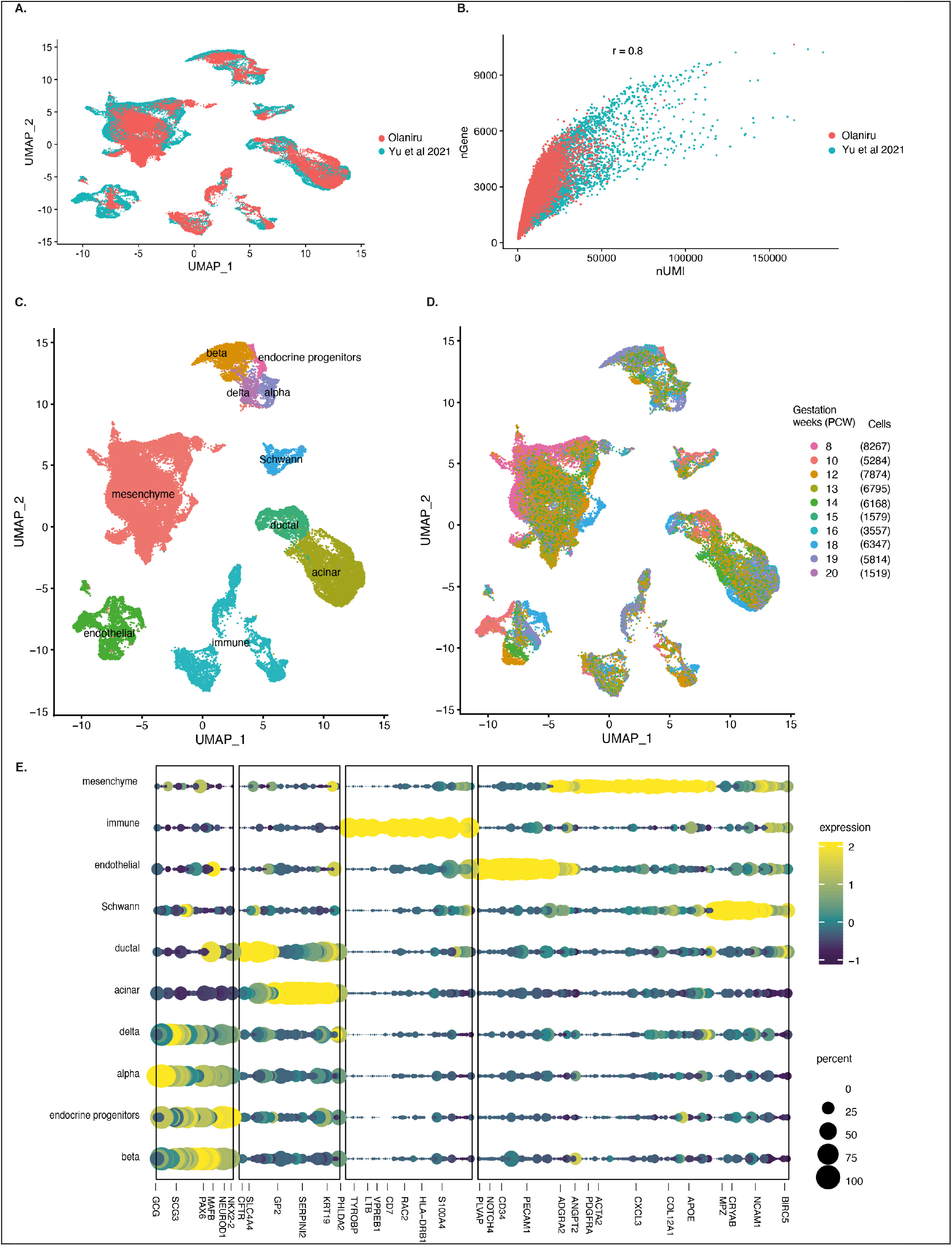
Analysis of the combined scRNA-seq data from this study and Yu et al., 2021. **A)** UMAP embedding showing integration of the two scRNA-seq datasets of the developing human pancreas at 8-20 PCW. **B)** Genes versus UMI plot showing a strong correlation between the merged scRNA-seq datasets. **C)** UMAP embedding showing the annotated cell clusters in the combined scRNA-seq data. **D)** UMAP embedding showing the different gestational stages and the number of cells per stage. **E)** Stacked violin plot showing differentially expressed genes across the annotated clusters.

**Supplementary Figure 5:**
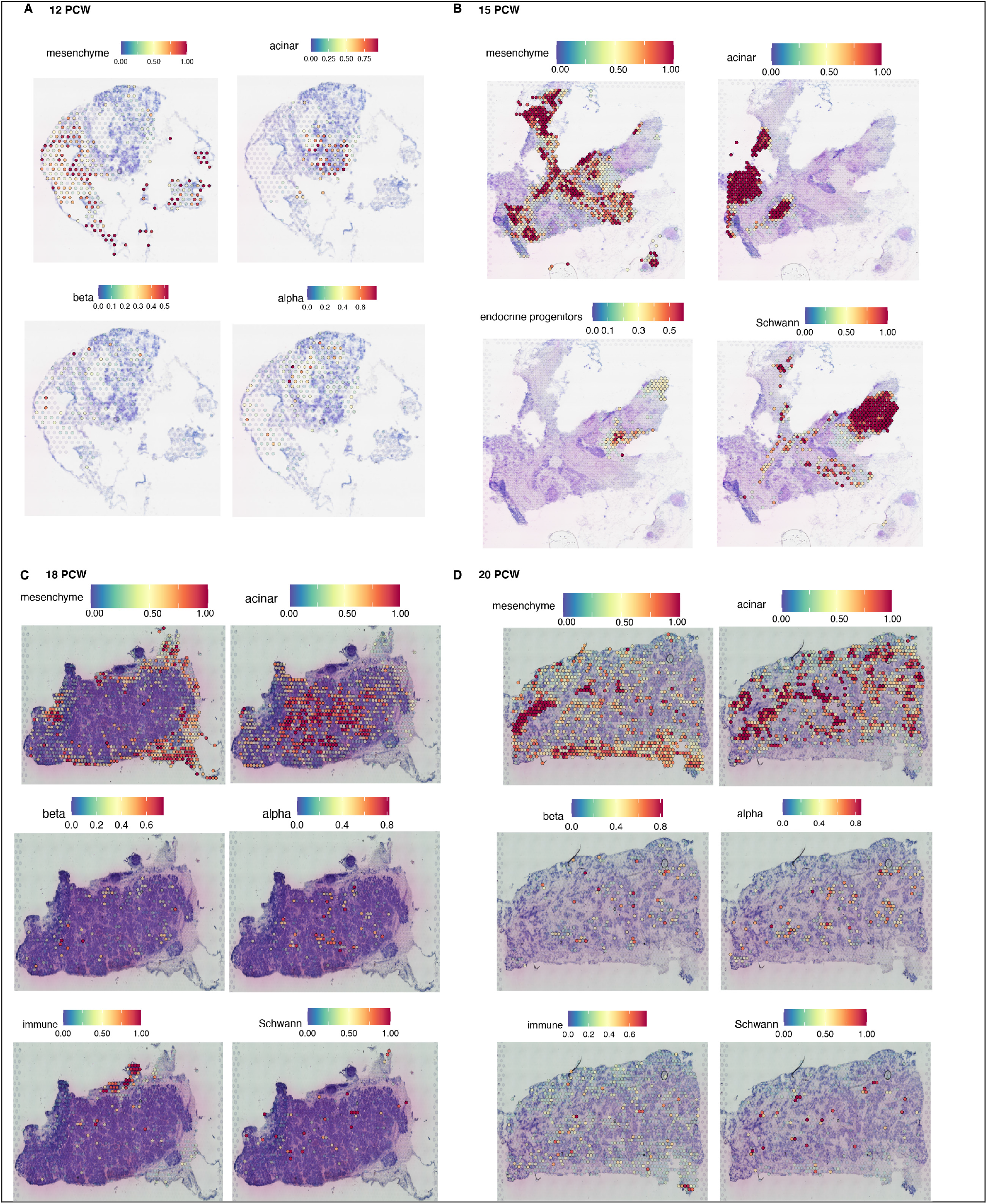
Cell type prediction of spatial samples using cell annotations from the combined scRNA-seq data. **A)** Predicted localisation of mesenchymal, acinar, beta and alpha cells in a 12 PCW pancreas. **B)** Predicted localisation of mesenchymal, acinar, endocrine progenitors and Schwann cells at 15 PCW. **C)** Predicted localisation of mesenchymal, acinar, beta, alpha, immune and Schwann cell populations at 18 PCW. **D)** Predicted localisation of mesenchymal, acinar, beta, alpha, immune and Schwann cells at 20 PCW.

**Supplementary Figure 6:**
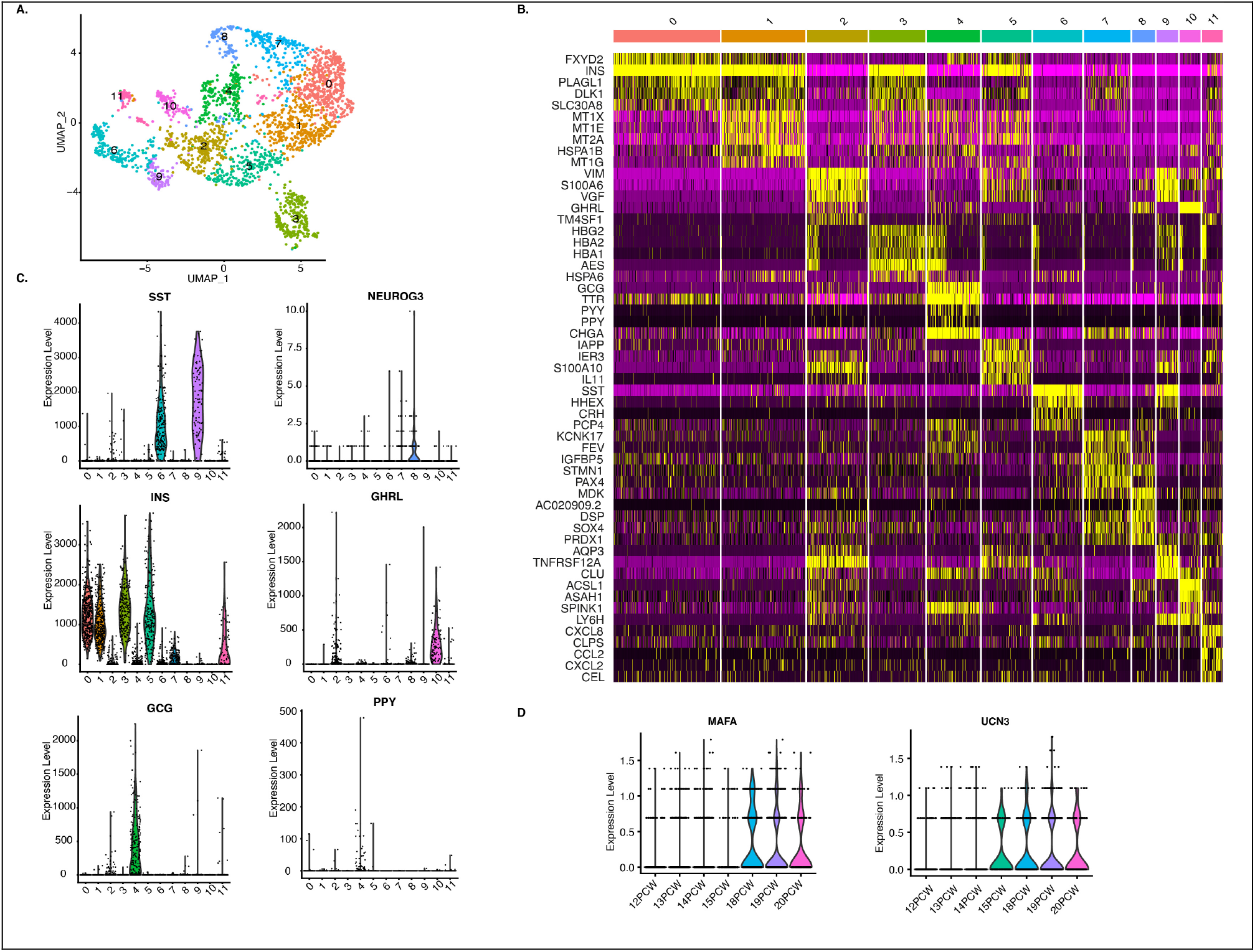
Identification of cell types within the endocrine compartment. **A)** UMAP embedding showing 11 subclusters from the reclustering of the endocrine cells. **B)** Heatmap showing differentially expressed genes in the 11 endocrine subclusters. **C)** Violin plots showing expression of key islet genes within each subcluster. **D)** Violin plot showing expression of the beta cell maturity markers MAFA and UNC3 at different gestational stages.

**Supplementary Figure 7:**
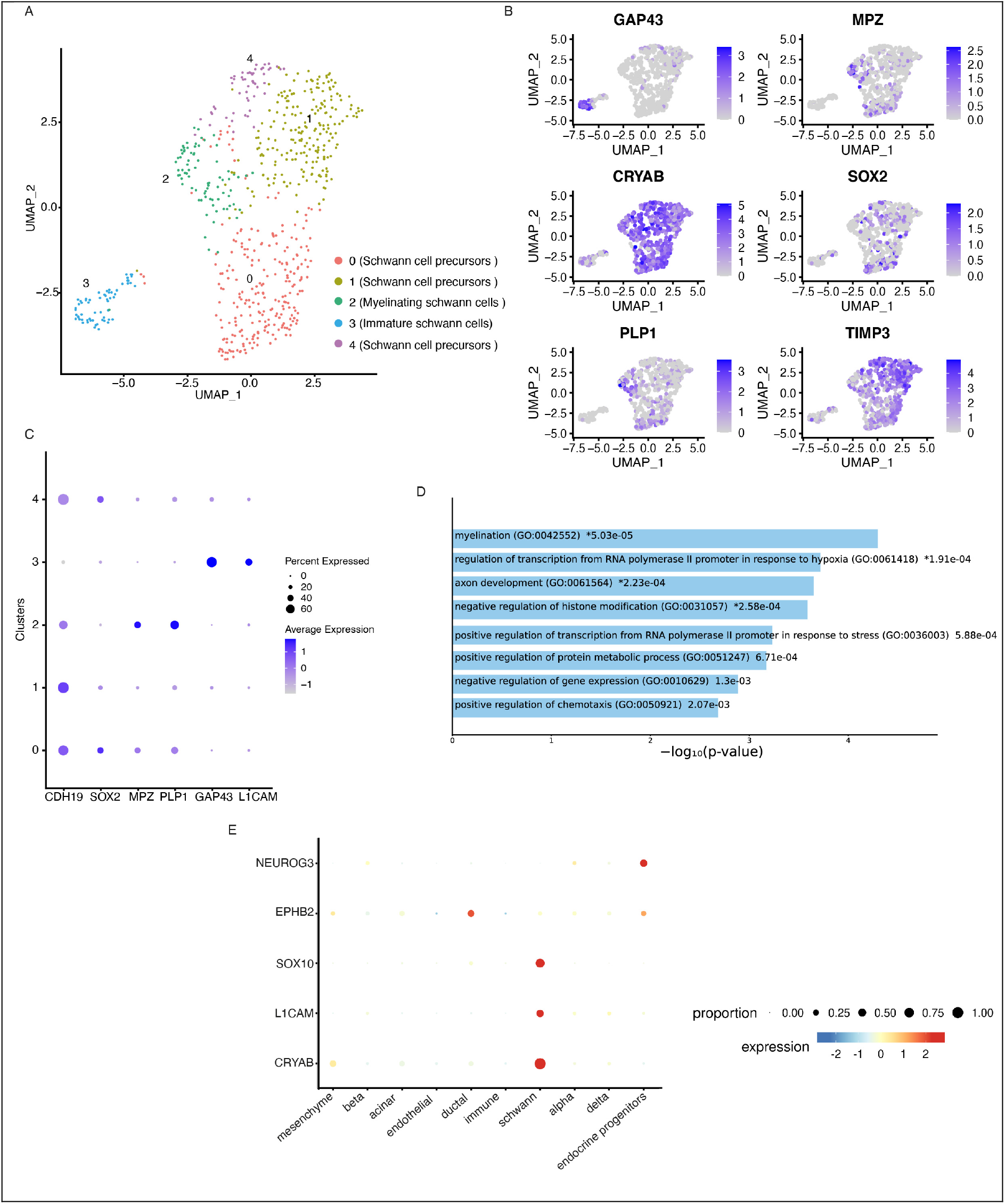
Identification of Schwann cell populations in developing human pancreas at 12-20 PCW. **A)** UMAP embedding showing five Schwann cell subclusters. **B)** Feature plots showing expression of select Schwann markers. **C)** Dot plot depicting marker genes used to identify Schwann cell subclusters. **D)** Gene ontology analysis showing biological functions regulated by genes that are more than 3-fold differentially expressed in Schwann cell subcluster 2. **E)** Dot plot showing that L1CAM expression is specific to the Schwann cell populations while EPHB2 is expressed by ductal and endocrine progenitors.

**Supplementary Figure 8:**
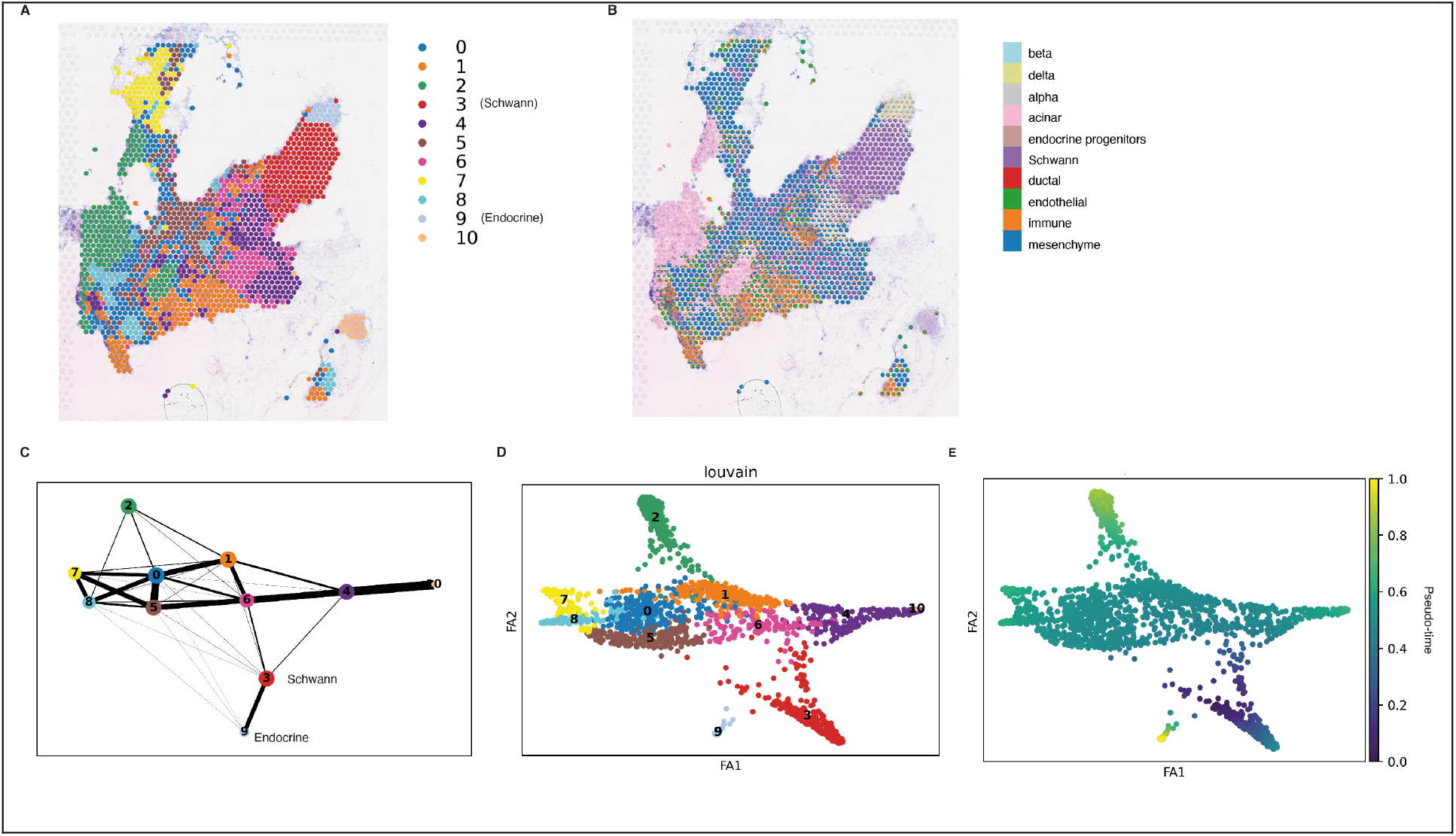
Global trajectory analysis at 15 PCW. **A)** stLearn clustering revealed 10 cell clusters at 15 PCW. Clusters 3 and 9 were annotated as Schwann and endocrine populations, respectively. **B)** Cell type predictions inferred from scRNA-seq data showing the different cell types at 15 PCW. **C)** PAGA trajectory inference plot showing connections between the clusters. **D)** Clustering and **E)** diffusion pseudotime places Cluster 3 (Schwann cells) as the root of the trajectory since it is earliest in the computed pseudo-time.

## Notes

### Competing Interest Statement

The authors have declared no competing interest.

## REFERENCES

Ariza, L., Cañete, A., Rojas, A., Muñoz-Chápuli, R., and Carmona, R. (2018). Role of the Wilms’ tumor suppressor gene Wt1 in pancreatic development. Dev. Dyn. 247, 924–933.

Armstrong, J.F., Pritchard-Jones, K., Bickmore, W.A., Hastie, N.D., and Bard, J.B. (1993). The expression of the Wilms’ tumour gene, WT1, in the developing mammalian embryo. Mech. Dev. 40, 85–97.

Arnes, L., Hill, J.T., Gross, S., Magnuson, M.A., and Sussel, L. (2012). Ghrelin expression in the mouse pancreas defines a unique multipotent progenitor population. PLoS One 7, e52026.

Becht, E., McInnes, L., Healy, J., Dutertre, C.-A., Kwok, I.W.H., Ng, L.G., Ginhoux, F., and Newell, E.W. (2018). Dimensionality reduction for visualizing single-cell data using UMAP. Nat. Biotechnol. 37, 38–44.

Bray, F., Ferlay, J., Soerjomataram, I., Siegel, R.L., Torre, L.A., and Jemal, A. (2018). Global cancer statistics 2018: GLOBOCAN estimates of incidence and mortality worldwide for 36 cancers in 185 countries. CA Cancer J Clin 68, 394–424.

Byrnes, L.E., Wong, D.M., Subramaniam, M., Meyer, N.P., Gilchrist, C.L., Knox, S.M., Tward, A.D., Ye, C.J., and Sneddon, J.B. (2018). Lineage dynamics of murine pancreatic development at single-cell resolution. Nat. Commun. 9, 3922.

Cabrera, O., Berman, D.M., Kenyon, N.S., Ricordi, C., Berggren, P.-O., and Caicedo, A. (2006). The unique cytoarchitecture of human pancreatic islets has implications for islet cell function. Proc. Natl. Acad. Sci. USA 103, 2334–2339.

Cao, J., Spielmann, M., Qiu, X., Huang, X., Ibrahim, D.M., Hill, A.J., Zhang, F., Mundlos, S., Christiansen, L., Steemers, F.J., et al. (2019). The single-cell transcriptional landscape of mammalian organogenesis. Nature 566, 496–502.

Cogger, K.F., Sinha, A., Sarangi, F., McGaugh, E.C., Saunders, D., Dorrell, C., Mejia-Guerrero, S., Aghazadeh, Y., Rourke, J.L., Screaton, R.A., et al. (2017). Glycoprotein 2 is a specific cell surface marker of human pancreatic progenitors. Nat. Commun. 8, 331.

D’Amour, K.A., Bang, A.G., Eliazer, S., Kelly, O.G., Agulnick, A.D., Smart, N.G., Moorman, M.A., Kroon, E., Carpenter, M.K., and Baetge, E.E. (2006). Production of pancreatic hormone-expressing endocrine cells from human embryonic stem cells. Nat. Biotechnol. 24, 1392–1401.

Dries, R., Zhu, Q., Dong, R., Eng, C.-H.L., Li, H., Liu, K., Fu, Y., Zhao, T., Sarkar, A., Bao, F., et al. (2021). Giotto: a toolbox for integrative analysis and visualization of spatial expression data. Genome Biol. 22, 78.

Efremova, M., Vento-Tormo, M., Teichmann, S.A., and Vento-Tormo, R. (2020). CellPhoneDB: inferring cell-cell communication from combined expression of multi-subunit ligand-receptor complexes. Nat. Protoc. 15, 1484–1506.

Emmanouilidou, A., Karetsou, Z., Tzima, E., Kobayashi, T., and Papamarcaki, T. (2013). Knockdown of prothymosin α leads to apoptosis and developmental defects in zebrafish embryos. Biochem Cell Biol 91, 325–332.

Fujino, A., Pieretti-Vanmarcke, R., Wong, A., Donahoe, P.K., and Arango, N.A. (2007). Sexual dimorphism of G-protein subunit Gng13 expression in the cortical region of the developing mouse ovary. Dev. Dyn. 236, 1991–1996.

Garcia-Alonso, L., Handfield, L.-F., Roberts, K., Nikolakopoulou, K., Fernando, R.C., Gardner, L., Woodhams, B., Arutyunyan, A., Polanski, K., Hoo, R., et al. (2021). Mapping the temporal and spatial dynamics of the human endometrium in vivo and in vitro. Nat. Genet.

Gittes, G.K., Galante, P.E., Hanahan, D., Rutter, W.J., and Debase, H.T. (1996). Lineage-specific morphogenesis in the developing pancreas: role of mesenchymal factors. Development 122, 439– 447.

Gonçalves, C.A., Larsen, M., Jung, S., Stratmann, J., Nakamura, A., Leuschner, M., Hersemann, L., Keshara, R., Perlman, S., Lundvall, L., et al. (2021). A 3D system to model human pancreas development and its reference single-cell transcriptome atlas identify signaling pathways required for progenitor expansion. Nat. Commun. 12, 3144.

Gonda, Y., Sakurai, H., Hirata, Y., Tabata, H., Ajioka, I., and Nakajima, K. (2007). Expression profiles of Insulin-like growth factor binding protein-like 1 in the developing mouse forebrain. Gene Expr Patterns 7, 431–440.

Hafemeister, C., and Satija, R. (2019). Normalization and variance stabilization of single-cell RNA-seq data using regularized negative binomial regression. Genome Biol. 20, 296.

Haghverdi, L., Büttner, M., Wolf, F.A., Buettner, F., and Theis, F.J. (2016). Diffusion pseudotime robustly reconstructs lineage branching. Nat. Methods 13, 845–848.

Hänzelmann, S., Castelo, R., and Guinney, J. (2013). GSVA: gene set variation analysis for microarray and RNA-seq data. BMC Bioinformatics 14, 7.

Hibsher, D., Epshtein, A., Oren, N., and Landsman, L. (2016). Pancreatic Mesenchyme Regulates Islet Cellular Composition in a Patched/Hedgehog-Dependent Manner. Sci. Rep. 6, 38008.

Huang, Y., McCarthy, D.J., and Stegle, O. (2019). Vireo: Bayesian demultiplexing of pooled single-cell RNA-seq data without genotype reference. Genome Biol. 20, 273.

Jessen, K.R., and Mirsky, R. (2005). The origin and development of glial cells in peripheral nerves. Nat. Rev. Neurosci. 6, 671–682.

Kriegebaum, C.B., Gutknecht, L., Bartke, L., Reif, A., Buttenschon, H.N., Mors, O., Lesch, K.-P., and Schmitt, A.G. (2010). The expression of the transcription factor FEV in adult human brain and its association with affective disorders. J. Neural Transm. 117, 831–836.

Kuleshov, M.V., Jones, M.R., Rouillard, A.D., Fernandez, N.F., Duan, Q., Wang, Z., Koplev, S., Jenkins, S.L., Jagodnik, K.M., Lachmann, A., et al. (2016). Enrichr: a comprehensive gene set enrichment analysis web server 2016 update. Nucleic Acids Res. 44, W90–7.

Landsman, L., Nijagal, A., Whitchurch, T.J., Vanderlaan, R.L., Zimmer, W.E., Mackenzie, T.C., and Hebrok, M. (2011). Pancreatic mesenchyme regulates epithelial organogenesis throughout development. PLoS Biol. 9, e1001143.

Li, L., Miano, J.M., Cserjesi, P., and Olson, E.N. (1996). SM22 alpha, a marker of adult smooth muscle, is expressed in multiple myogenic lineages during embryogenesis. Circ. Res. 78, 188–195.

Liu, Z., Jin, Y.-Q., Chen, L., Wang, Y., Yang, X., Cheng, J., Wu, W., Qi, Z., and Shen, Z. (2015). Specific marker expression and cell state of Schwann cells during culture in vitro. PLoS One 10, e0123278.

Majesky, M.W., Dong, X.R., Regan, J.N., and Hoglund, V.J. (2011). Vascular smooth muscle progenitor cells: building and repairing blood vessels. Circ. Res. 108, 365–377.

Mantri, M., Scuderi, G.J., Abedini-Nassab, R., Wang, M.F.Z., McKellar, D., Shi, H., Grodner, B., Butcher, J.T., and De Vlaminck, I. (2021). Spatiotemporal single-cell RNA sequencing of developing chicken hearts identifies interplay between cellular differentiation and morphogenesis. Nat. Commun. 12, 1771.

Murtaugh, L.C. (2008). The what, where, when and how of Wnt/β-catenin signaling in pancreas development. Organogenesis 4, 81–86.

Nair, G., and Hebrok, M. (2015). Islet formation in mice and men: lessons for the generation of functional insulin-producing β-cells from human pluripotent stem cells. Curr. Opin. Genet. Dev. 32, 171–180.

Nair, G.G., Liu, J.S., Russ, H.A., Tran, S., Saxton, M.S., Chen, R., Juang, C., Li, M.-L., Nguyen, V.Q., Giacometti, S., et al. (2019). Recapitulating endocrine cell clustering in culture promotes maturation of human stem-cell-derived β cells. Nat. Cell Biol. 21, 263–274.

Namvar, S., Woolf, A.S., Zeef, L.A., Wilm, T., Wilm, B., and Herrick, S.E. (2018). Functional molecules in mesothelial-to-mesenchymal transition revealed by transcriptome analyses. J. Pathol. 245, 491–501.

Olerud, J., Kanaykina, N., Vasylovska, S., King, D., Sandberg, M., Jansson, L., and Kozlova, E.N. (2009). Neural crest stem cells increase beta cell proliferation and improve islet function in co-transplanted murine pancreatic islets. Diabetologia 52, 2594–2601.

Ouyang, G., Pan, G., Liu, Q., Wu, Y., Liu, Z., Lu, W., Li, S., Zhou, Z., and Wen, Y. (2020). The global, regional, and national burden of pancreatitis in 195 countries and territories, 1990-2017: a systematic analysis for the Global Burden of Disease Study 2017. BMC Med. 18, 388.

Pagliuca, F.W., Millman, J.R., Gürtler, M., Segel, M., Van Dervort, A., Ryu, J.H., Peterson, Q.P., Greiner, D., and Melton, D.A. (2014). Generation of functional human pancreatic β cells in vitro. Cell 159, 428–439.

Pearse, A.G., and Polak, J.M. (1971). Neural crest origin of the endocrine polypeptide (APUD) cells of the gastrointestinal tract and pancreas. Gut 12, 783–788.

Perera, S.N., and Kerosuo, L. (2021). On the road again: Establishment and maintenance of stemness in the neural crest from embryo to adulthood. Stem Cells 39, 7–25.

Pham, D.T., Tan, X., Xu, J., Grice, L.F., Lam, P.Y., Raghubar, A., Vukovic, J., Ruitenberg, M.J., and Nguyen, Q.H. (2020). stLearn: integrating spatial location, tissue morphology and gene expression to find cell types, cell-cell interactions and spatial trajectories within undissociated tissues. BioRxiv.

Pictet, R.L., Rall, L.B., Phelps, P., and Rutter, W.J. (1976). The neural crest and the origin of the insulin-producing and other gastrointestinal hormone-producing cells. Science 191, 191–192.

Que, J., Wilm, B., Hasegawa, H., Wang, F., Bader, D., and Hogan, B.L.M. (2008). Mesothelium contributes to vascular smooth muscle and mesenchyme during lung development. Proc. Natl. Acad. Sci. USA 105, 16626–16630.

Ramond, C., Beydag-Tasöz, B.S., Azad, A., van de Bunt, M., Petersen, M.B.K., Beer, N.L., Glaser, N., Berthault, C., Gloyn, A.L., Hansson, M., et al. (2018). Understanding human fetal pancreas development using subpopulation sorting, RNA sequencing and single-cell profiling. Development 145.

Raredon, M.S.B., Yang, J., Garritano, J., Wang, M., Kushnir, D., Schupp, J.C., Adams, T.S., Greaney, A.M., Leiby, K.L., Kaminski, N., et al. (2021). Connectome : computation and visualization of cell-cell signaling topologies in single-cell systems data. BioRxiv.

Rezania, A., Bruin, J.E., Arora, P., Rubin, A., Batushansky, I., Asadi, A., O’Dwyer, S., Quiskamp, N., Mojibian, M., Albrecht, T., et al. (2014). Reversal of diabetes with insulin-producing cells derived in vitro from human pluripotent stem cells. Nat. Biotechnol. 32, 1121–1133.

Russ, H.A., Parent, A.V., Ringler, J.J., Hennings, T.G., Nair, G.G., Shveygert, M., Guo, T., Puri, S., Haataja, L., Cirulli, V., et al. (2015). Controlled induction of human pancreatic progenitors produces functional beta-like cells in vitro. EMBO J. 34, 1759–1772.

Saeedi, P., Petersohn, I., Salpea, P., Malanda, B., Karuranga, S., Unwin, N., Colagiuri, S., Guariguata, L., Motala, A.A., Ogurtsova, K., et al. (2019). Global and regional diabetes prevalence estimates for 2019 and projections for 2030 and 2045: Results from the International Diabetes Federation Diabetes Atlas, 9th edition. Diabetes Res. Clin. Pract. 157, 107843.

Salinno, C., Cota, P., Bastidas-Ponce, A., Tarquis-Medina, M., Lickert, H., and Bakhti, M. (2019). β-Cell Maturation and Identity in Health and Disease. Int. J. Mol. Sci. 20.

Schaefer, T., and Lengerke, C. (2020). SOX2 protein biochemistry in stemness, reprogramming, and cancer: the PI3K/AKT/SOX2 axis and beyond. Oncogene 39, 278–292.

Seymour, P.A., Freude, K.K., Tran, M.N., Mayes, E.E., Jensen, J., Kist, R., Scherer, G., and Sander, M. (2007). SOX9 is required for maintenance of the pancreatic progenitor cell pool. Proc. Natl. Acad. Sci. USA 104, 1865–1870.

Stoeckius, M., Zheng, S., Houck-Loomis, B., Hao, S., Yeung, B.Z., Mauck, W.M., Smibert, P., and Satija, R. (2018). Cell Hashing with barcoded antibodies enables multiplexing and doublet detection for single cell genomics. Genome Biol. 19, 224.

Stuart, T., Butler, A., Hoffman, P., Hafemeister, C., Papalexi, E., Mauck, W.M., Hao, Y., Stoeckius, M., Smibert, P., and Satija, R. (2019). Comprehensive Integration of Single-Cell Data. Cell 177, 1888–1902.e21.

Su, Y., Jono, H., Misumi, Y., Senokuchi, T., Guo, J., Ueda, M., Shinriki, S., Tasaki, M., Shono, M., Obayashi, K., et al. (2012). Novel function of transthyretin in pancreatic alpha cells. FEBS Lett. 586, 4215–4222.

Takahashi, M., and Osumi, N. (2005). Identification of a novel type II classical cadherin: rat cadherin19 is expressed in the cranial ganglia and Schwann cell precursors during development. Dev. Dyn. 232, 200–208.

Tran, T.N., and Bader, G.D. (2020). Tempora: Cell trajectory inference using time-series single-cell RNA sequencing data. PLoS Comput. Biol. 16, e1008205.

Trapnell, C., Cacchiarelli, D., Grimsby, J., Pokharel, P., Li, S., Morse, M., Lennon, N.J., Livak, K.J., Mikkelsen, T.S., and Rinn, J.L. (2014). The dynamics and regulators of cell fate decisions are revealed by pseudotemporal ordering of single cells. Nat. Biotechnol. 32, 381–386.

van der Meulen, T., Xie, R., Kelly, O.G., Vale, W.W., Sander, M., and Huising, M.O. (2012). Urocortin 3 marks mature human primary and embryonic stem cell-derived pancreatic alpha and beta cells. PLoS One 7, e52181.

Veres, A., Faust, A.L., Bushnell, H.L., Engquist, E.N., Kenty, J.H.-R., Harb, G., Poh, Y.-C., Sintov, E., Gürtler, M., Pagliuca, F.W., et al. (2019). Charting cellular identity during human in vitro β-cell differentiation. Nature 569, 368–373.

Villasenor, A., Chong, D.C., and Cleaver, O. (2008). Biphasic Ngn3 expression in the developing pancreas. Dev. Dyn. 237, 3270–3279.

Wells, J.M., Esni, F., Boivin, G.P., Aronow, B.J., Stuart, W., Combs, C., Sklenka, A., Leach, S.D., and Lowy, A.M. (2007). Wnt/beta-catenin signaling is required for development of the exocrine pancreas. BMC Dev. Biol. 7, 4.

Wilm, B., Ipenberg, A., Hastie, N.D., Burch, J.B.E., and Bader, D.M. (2005). The serosal mesothelium is a major source of smooth muscle cells of the gut vasculature. Development 132, 5317–5328.

Wolf, F.A., Hamey, F.K., Plass, M., Solana, J., Dahlin, J.S., Göttgens, B., Rajewsky, N., Simon, L., and Theis, F.J. (2019). PAGA: graph abstraction reconciles clustering with trajectory inference through a topology preserving map of single cells. Genome Biol. 20, 59.

Yu, X.-X., Qiu, W.-L., Yang, L., Wang, Y.-C., He, M.-Y., Wang, D., Zhang, Y., Li, L.-C., Zhang, J., Wang, Y., et al. (2021). Sequential progenitor states mark the generation of pancreatic endocrine lineages in mice and humans. Cell Res. 31, 886–903.

Yue, R., Shen, B., and Morrison, S.J. (2016). Clec11a/osteolectin is an osteogenic growth factor that promotes the maintenance of the adult skeleton. Elife 5.

Zhu, Q., Song, L., Peng, G., Sun, N., Chen, J., Zhang, T., Sheng, N., Tang, W., Qian, C., Qiao, Y., et al. (2014). The transcription factor Pou3f1 promotes neural fate commitment via activation of neural lineage genes and inhibition of external signaling pathways. Elife 3.

Zhu, S., Russ, H.A., Wang, X., Zhang, M., Ma, T., Xu, T., Tang, S., Hebrok, M., and Ding, S. (2016). Human pancreatic beta-like cells converted from fibroblasts. Nat. Commun. 7, 10080.

